# Distinct chromosomal “niches” in the genome of *S. cerevisiae* provide the background for genomic innovation and shape the fate of gene duplicates

**DOI:** 10.1101/2022.02.26.482092

**Authors:** Athanasia Stavropoulou, Aimilios Tassios, Maria Kalyva, Michalis Georgoulopoulos, Nikolaos Vakirlis, Ioannis Iliopoulos, Christoforos Nikolaou

## Abstract

Gene duplication is a major source of genomic innovation in all eukaryotes, with large proportions of genes being the result of either small-scale (SSD) or genome-wide duplication (WGD) events. In the model unicellular eukaryote *Saccharomyces cerevisiae*, of which nearly one third of the genome corresponds to gene duplicates, the two modes of duplication have been shown to follow different evolutionary fates, with SSD genes being more prone to acquire novel functionalities (neofunctionalization) and WGD more likely to retain different parts of the original ancestral function (subfunctionalization). Having previously described aspects of functional compartmentalization for the genes of *S. cerevisiae*, in this work we set out to investigate the existence of positional preferences of gene duplicates. We found that SSD and WGD genes are organized in distinct gene clusters that are, furthermore, segregated, occupying different regions, with SSD being more peripheral and WGD more centrally positioned close to centromeric chromatin.

Duplicate gene clusters differ from the rest of the genome in terms of gene size and spacing, gene expression variability and regulatory complexity. What is more interesting, some of these properties are also shared by singleton genes residing in duplicate-rich regions in a position-dependent manner. Our analysis further reveals particular chromatin architectures in the promoters of duplicate genes, which are generally longer, with less pronounced nucleosome-free regions, strong structural constraints and a larger number of regulatory elements. Such structural features appear to be important for gene evolution as we find SSD gene-pair co-expression to be strongly associated with the similarity of nucleosome positioning patterns.

We propose that specific regions of the yeast genome provide a favourable environment for the generation and maintenance of small-scale gene duplicates. The existence of such genomic “niches” is supported by the enrichment of these regions in singleton genes bearing similarities with gene “relics”, remnants of recent duplications that have reverted to single gene status. Our findings provide a valuable framework for the study of genomic innovation and suggest taking into account positional preferences in the study of gene emergence and fixation in experimentally and naturally evolving populations.

## Introduction

Gene duplication is widely accepted to be a major source of genomic innovation, occurring with relatively high frequency, through various mechanisms and affecting large proportions of chromosomes or even entire genomes (Ohno 1970; Acharya and Ghosh 2016). Duplicated genes may be the result of either localized, short-scale duplication (SSD), through replication slippage or ectopic recombination (Zhang 2003), or of cataclysmic events of whole-genome duplication (WGD) leading to temporary polyploidy (Otto and Whitton 2000; Sémon and Wolfe 2007). Both phenomena are more common than one might expect (Graur and Li 2000; Lynch and Conery 2003) but genomes are not replete with multiple copies of genes. This is because once a gene is duplicated, it becomes subject to strong evolutionary constraints that vary in both type and extent. Functional redundancy and the energy cost of maintaining two identical genes leads to the most likely outcome, which is the subsequent erosion and eventual loss of one of the two copies, through the accumulation of deleterious mutations (Dujon et al. 2004; Van Hoek and Hogeweg 2007; Innan and Kondrashov 2010). In some cases, however, duplicated genes may diverge to the point of acquiring diversified functions, which leads to the fixation of both copies. This process also varies, depending on whether the two copies both diverge to assume subsets of the original functions of the ancestral gene (sub-functionalization) or if one adopts a novel role, while the other preserves the original (neo-functionalization) (He and Zhang 2005; Sandve et al. 2018).

The way genomic and other constraints drive the fate of gene duplicates has been investigated in great depth in the case of the budding yeast *Saccharomyces cerevisiae*. Nearly one third of *S. cerevisiae* genes are the result of gene duplication, roughly half of which originate from a massive whole-genome duplication event that occurred ∼100 million years ago (Wolfe and Shields 1997; Kellis et al. 2004; Wolfe 2015), while the rest are the result of continuously occurring, small-scale duplications. A number of properties appear to be particular to yeast duplicated genes in general. Since relatively early, they were shown to be over-represented among highly expressed genes (Seoighe and Wolfe 1999), to reside in early replicating chromatin (Guan et al. 2007), to have more complex promoters (Papp et al. 2003; Wapinski et al. 2007) and, overall, fewer genetic interactions than non-duplicated, singleton genes (Costanzo et al. 2010).

The gene duplication mechanism may also be associated with different properties between “ohnologues”, that is whole-genome (WGD) and paralogs, which are small-scale duplicate (SSD) (Baudot et al. 2004). Knock-out studies showed WGD genes to be less essential and genetic interaction profiling suggested WGD pairs to be functionally more similar than SSD ones (Hakes, Pinney, et al. 2007) WGD are aslo more likely to form part of protein complexes (Hakes, Lovell, et al. 2007). SSD genes, on the other hand were found to have greater expression divergence (Papp et al. 2003), to be more prone to accumulate non-synonymous substitutions (Keane et al. 2014), as well as new protein-protein interactions (Fares et al. 2013). Together these observations draw associations with the evolutionary fate of gene duplicates, with WGD being more likely to be maintained due to subfunctionalization and the maintenance of dosage balance, while SSD through the adoption of divergent expression patterns and the acquisition of novel functions (Mattenberger et al. 2017).

Up to date, most of the analyses have treated the two daughter genes as linked or entangled from their birth through an “invisible thread”, regardless of their genomic context. Nevertheless the genomic environment, in which a gene is embedded, is an important determinant of its function and evolution, shaped by epigenetic factors and access by transcriptional regulators in the three-dimensional genomic landscape (De and Babu 2010; Tsochatzidou et al. 2017). Beyond sequence and functional conservation, the spatial organization of yeast genes has been the subject of extensive research. The conservation of pre- and post-WGD gene arrangement has been thoroughly documented by (Byrne and Wolfe 2005). The order of genes in linear chromosomes has been associated with the length of their intergenic spacers (Poyatos and Hurst 2007) as has the directionality of their transcription (Sugino and Innan 2012), while (Byrnes et al. 2006) have suggested that large intergenic spacers may decouple the transcriptional interference between adjacent genes and thus increase their expression divergence. In the context of gene duplication evolution (Fares et al. 2017) showed a strong tendency for WGD (but not SSD) to be located in genomic areas with increased non-synonymous substitution rate (mutational hotspots). WGD genes are also documented to very rarely undergo small-scale duplication (Makino and McLysaght 2010). Experimental evidence for the evolutionary fate of gene duplication being context-dependent came from the approach of (Naseeb et al. 2017), in which a duplicated copy of the IFA38 gene was found to be more likely to escape deletion when positioned in tandem with the ancestral gene.

It is, thus, quite plausible that the position in which a gene is found and the overall genomic “context” may shape not only its regulatory and expression patterns, but also it evolutionary fate. Even more so, one may hypothesize, that particular areas of the genome may represent more or less “permissive” environments for the occurrence and maintenance of gene duplicates. Having previously identified positional preferences for genes associated with sequence composition (Papanikolaou et al. 2009) gene regulation (Nikolaou et al. 2013; Tsochatzidou et al. 2017) and chromatin structure (Nikolaou 2018), in this work we focus on such context-dependent properties for gene duplicates. Starting from the observation of extensive gene spatial clustering for both WGD and SSD genes, we go on to define genomic domains of WGD/SSD enrichment and to examine how the properties of these regions may be affecting not only the duplicate but also the singleton genes they harbour. We find SSD and WGD being largely segregated in distinct parts of the yeast genome. We further uncover a number of structural, regulatory and functional properties of gene duplicates that are domain-specific and which are, in addition, partially shared by neighbouring singleton genes. Our findings may support a model for the evolution of gene duplication events, according to which, the yeast genome may be divided in areas with differential capacity for genetic innovation, driven primarily by the divergence of the genes’ regulatory sequences. We find that SSD duplication preferentially occurs in confined areas of the yeast genome that constitute genomic “niches” favourable for faster divergence, thus enabling the emergence of novel functions.

## Methods

### Genomic coordinates in one and three dimensions

Chromosomal coordinates of yeast genes were obtained from UCSC using the sacCer2 (June 2008) assembly to achieve maximum compatibility with all the datasets used. SSD and WGD gene clusters were created by merging genomic regions containing two or more genes of the same type within a distance smaller than 10kb. Clusters containing both SSD and WGD genes corresponded less than 20% of the total and were assigned to the gene type that represented the majority in the cluster.

Distances to the centromeres and the chromosomal edges were calculated as ratios over the entire chromosomal length as described in (Tsochatzidou et al. 2017). Coding density was calculated as the percentage of coding sequences spanning a region of eleven genes, symmetrically flanking the gene in question.

For the three dimensional coordinates we used the published conformational model of the yeast genome (Duan et al. 2010) which has been resampled at gene resolution. We obtained gene positions by linearly interpolating the model’s control points to approximate the center base pair of each gene, which resulted in each gene being represented as a set of three coordinates in arbitrary space. Assuming the mean coordinates of all genes to correspond to the center of the genome in 3D space, we calculated its euclidean distance from each gene and then took quantiles of these distances to assign genes into three sections: Central, bottom quartile (lowest 25% of the distances), Intermediate, middle half (>25% and <75% of the distances) and Peripheral, top quartile (highest 25% of the distances).

### Enrichment analysis

Enrichments of genome coordinates and set overlaps were calculated as described in (Andreadis et al. 2014). The positional overlaps between gene coordinates were calculated with BedTools (Quinlan and Hall 2010) and the enrichment was defined as the observed-over-expected ratio of coordinate overlap. The expected value was calculated as the product of the two independent proportions of gene coordinates over the total genome size. Significance was assessed with a permutation test, by shuffling the smaller of the two coordinate sets while keeping the same number and size distribution of its segments. In all cases, one thousand such shuffles were performed. The reported p-values corresponded to the proportion of times a random shuffle yielded a value more extreme than the one observed.

### Gene Age

The phylogenetic age of *S. cerevisiae* genes was determined by phylostratigraphy. We performined BLASTP (States and Gish 1994) searches on all available Fungi proteomes in GenBank (794 unique species excluding *S. cerevisiae*, 1,266 total proteomes, downloaded December 2019) with an E-value cutoff of 1E-3. Age was defined by as the most recent common ancestor of species that shared a homologue. The NCBI Taxonomy common tree was used, resulting in genes classified in the following phylogenetic ages: species-specific, genus (*Saccharomyces*), family (saccharomycetaceae), order (saccharomycetales), division (ascomycota) or kingdom (fungi).

### Conservation, sequence divergence and structural constraint data

Sequence conservation was measured using phastCons (Siepel et al. 2005) precalculated scores for an alignment of seven *Saccharomyces* species. Divergence between duplicate gene pairs was obtained from (Fares et al. 2013) and was calculated at the level of amino acid sequence, taking into account the age of the duplicate (for SSD genes).

Structural constraints were assessed with the use of two models: the deep learning model of (Routhier et al. 2020) which attempts to capture the effect of nucleotide substitutions on nucleosome positioning and our own, simpler model based on the variability of nucleosome positioning predictive scores (Nikolaou et al. 2010).

Aggregation of phastCons and structural constraint values was done through the calculation of a mean score over the segment under question, thus taking the average of all single-nucleotide-resolution values over the complete length of each gene. Average gene profiles were created as vectors of binned averaged values for 1000-nucleotide regions symmetrically flanking each gene’s transcription start site, in bins of 10 nt.

### Nucleosome positioning and gene regulation

Nucleosome positioning data were obtained from a genome wide MNase profiling (Jiang and Pugh 2009). Average gene profiles were created as described above for the same regions and bin size. Nucleosome positioning similarities for duplicate gene pairs were calculated as Pearson correlation coefficients of the nucleosome positioning 10nt-bin profiles.

We used a dataset of highly reliable conserved transcription factor binding sites (MacIsaac et al. 2006) to assign a set of transcription factors (TF), to each gene in our duplicate dataset. Transcriptional regulation similarity between gene duplicate pairs was calculated as the Jaccard index of their corresponding TFs. The Jaccard similarity between two sets, is defined as the ratio of the size of their intersection over that of their union.

### Co-expression scores and transcriptional variability

We obtained normalized expression data from a compendium of ∼2400 experimental conditions from the SPELL database (Hibbs et al. 2007). We used SPELL’s pre-calculated Adjusted Co-expression Score (ACS) as a measure of gene co-expression. In order to assess expression variability, we calculated the standard deviation of gene expression levels for each gene across all conditions and then normalized it across genes with the use of a z-score.

### Protein complexity, protein-protein interactions and functional enrichment

We used the PFAM database (Mistry et al. 2021) API to assign the predicted functional protein domains to the protein sequences of the complete set of genes. For each protein we then calculated the proportion of its sequence being attributed to a PFAM domain as a proxy for protein complexity. Protein-protein interactions were obtained from the STRING database (Szklarczyk et al. 2021). Functional entanglement was assessed as the number of GO terms associated with each gene. Functional enrichments were assessed with the use of gProfiler (Reimand et al. 2007) and suitably adopted custom R functions.

### Association Rules

Association rules were extracted from the sets of genes against transcriptional regulators using the data by (MacIsaac et al. 2006) (see above). We employed the *arules* R package (Hahsler et al. 2005; Hahsler et al. 2022) through the Apriori algorithm and extracted the top 10% most significant associations in term of lift. Lift corresponds to the strength of the association between regulator and gene target, controling for the prevalence of the regulator in the dataset. The results were presented in the form of association networks between regulators.

## Results

### Duplicate genes are segregated in the yeast genome

Because of their mode of duplication, which predominantly takes place in tandem, small-scale duplicate (henceforth SSD) gene pairs tend to be found in close proximity to each other and are also more often located on the same chromosome (Figure 1A, 1B). As expected, SSD genes, when found on the same chromosome, are very often juxtaposed to each other (Figure 1C). On the other hand whole-genome duplicate (from now on WGD) genes are preferentially localized in regions that maintain synteny and which conserve the ancestral juxtaposition (Figure 1A). Both types of genes are therefore likely to cluster in linear genomic space. Our starting hypothesis was that clusters of SSD and WGD genes occupy distinct areas of the yeast genome. In order to quantify this observation, we created clusters of SSD and WGD genes by merging genomic regions containing genes of the same type within a distance of 10kb. One first indication for the strong clustering of duplicate genes was that the size distribution of the created, extended regions was smaller than the corresponding ones for random selections of equal numbers of genes (p<=0.001 for 1000 random permutations). The created SSD/WGD- clusters clearly occupy distinct genomic spaces in the yeast genome (Figure 1D).

**Figure 1A.**
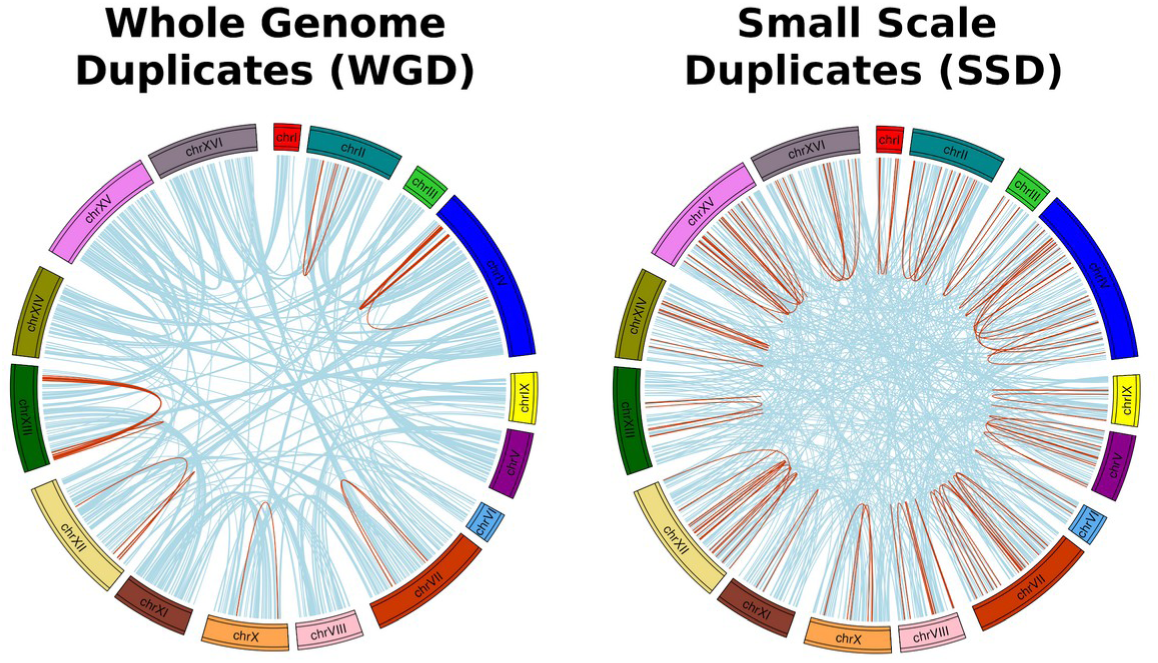
Circos Plots joining gene duplicate pairs in the yeast genome. Left: Whole Genome Duplicates (WGD), Right: Small-Scale Duplicates (SSD). Light blue lines join duplicate pairs on different chromosomes, while brown lines join pairs located on the same chromosome.

**Figure 1B.**
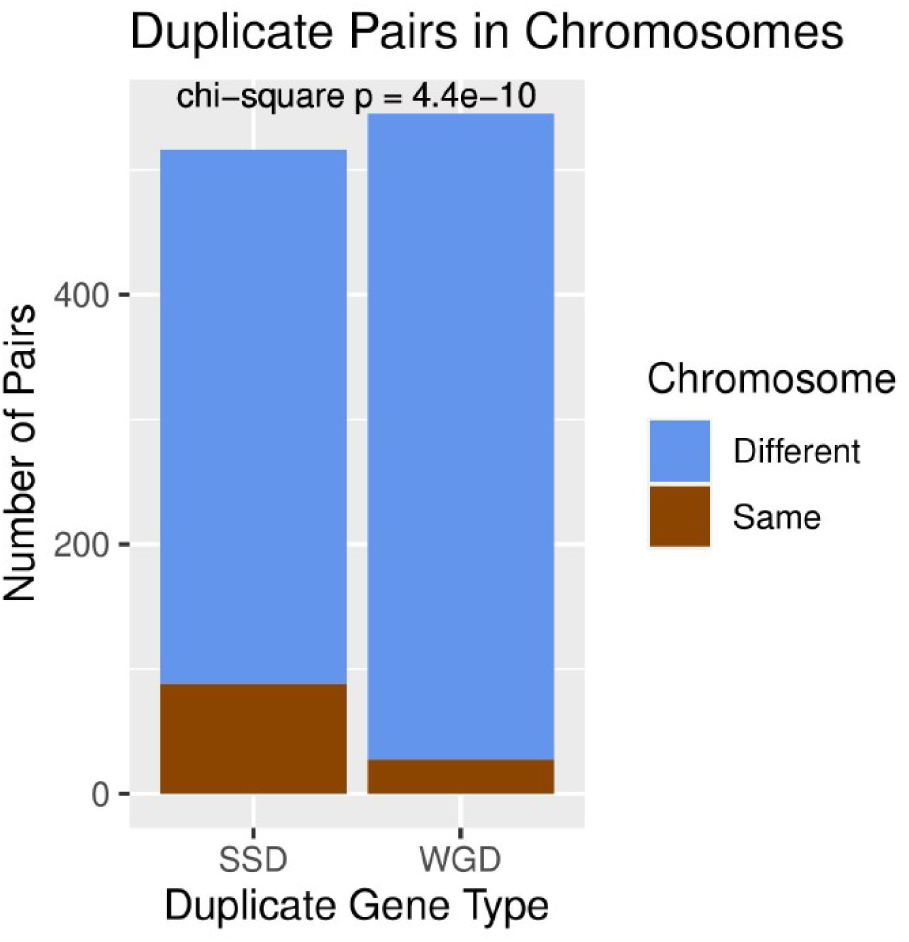
Number of gene duplicate pairs found on the same or on different chromosomes.

**Figure 1C.**
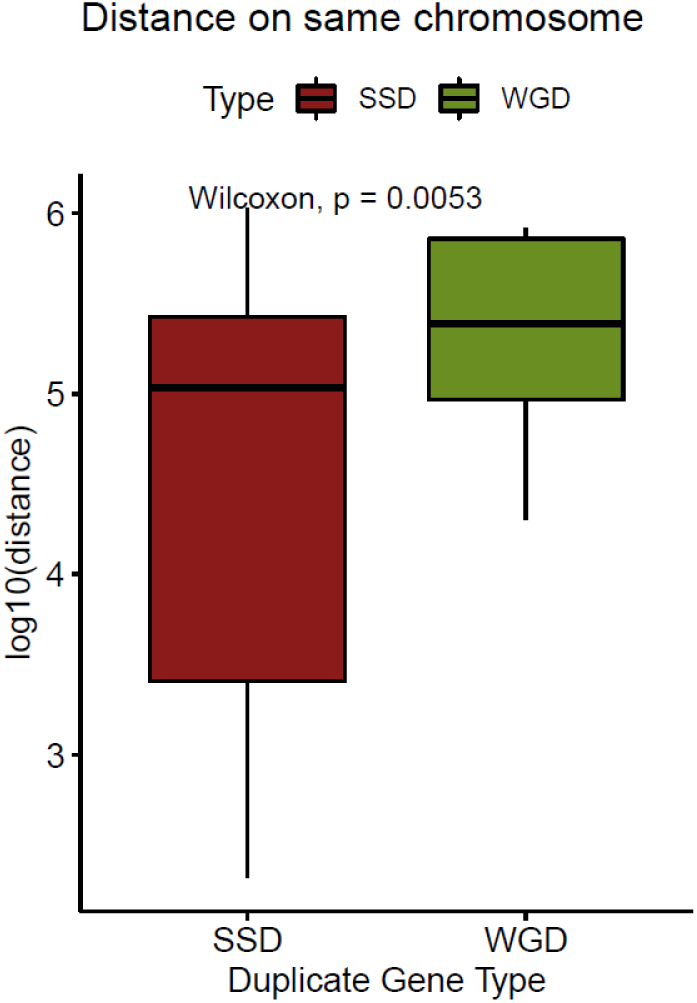
Distribution of linear distances (log10-transformed) between gene duplicate pairs that are located on the same chromosomes.

**Figure 1D.**
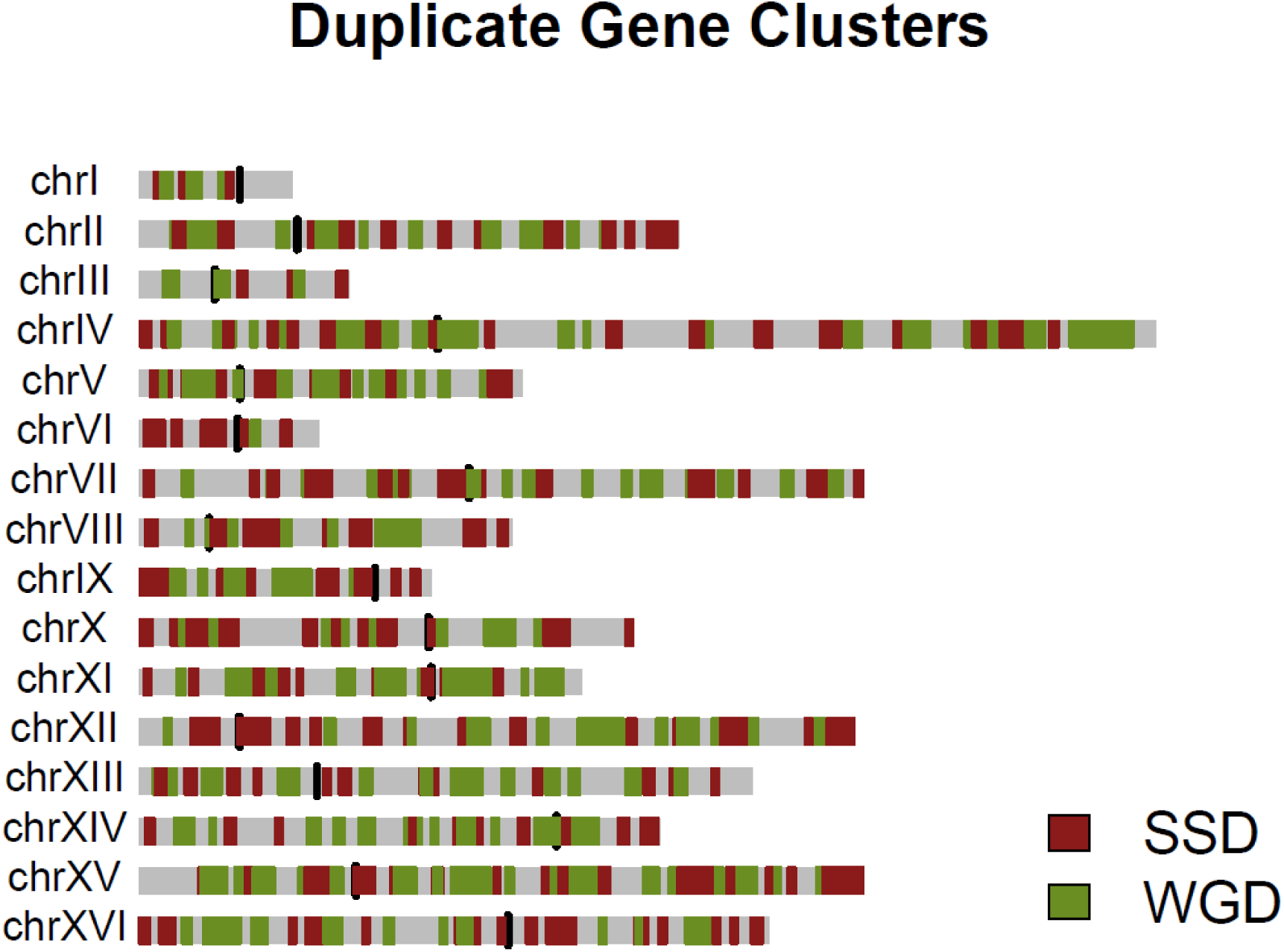
Genome domainogram showing the locations of gene duplicate clusters along the sixteen chromosomes of the yeast genome.

This is supported by a) the significantly small observed-over-expected overlap between the two types of clusters (o/e=0.7, p<0.001) and b) the distribution of relative distances obtained with the use of the *bedtools reldist* function (Quinlan and Hall 2010), which shows an enrichment towards relative distances that are considerably longer than the ones expected by chance (Figure 1E). The observed segregation of SSD and WGD genes is persistent for various cluster sizes, only to be lifted for cluster extension up to 100kb, which, by and large, correspond to large proportions of entire chromosomes. The localization of SSD (paralogs) and WGD (ohnologs) in different genomic areas has already been documented for vertebrates through the study of copy number variation (CNV), which may be largely responsible for the creation of SSD but to which WGD are largely refractory (Makino and McLysaght 2010; Makino et al. 2013). We here show that this is also the case for a much more coding dense genome such as the one of *S. cerevisiae*.

**Figure 1E.**
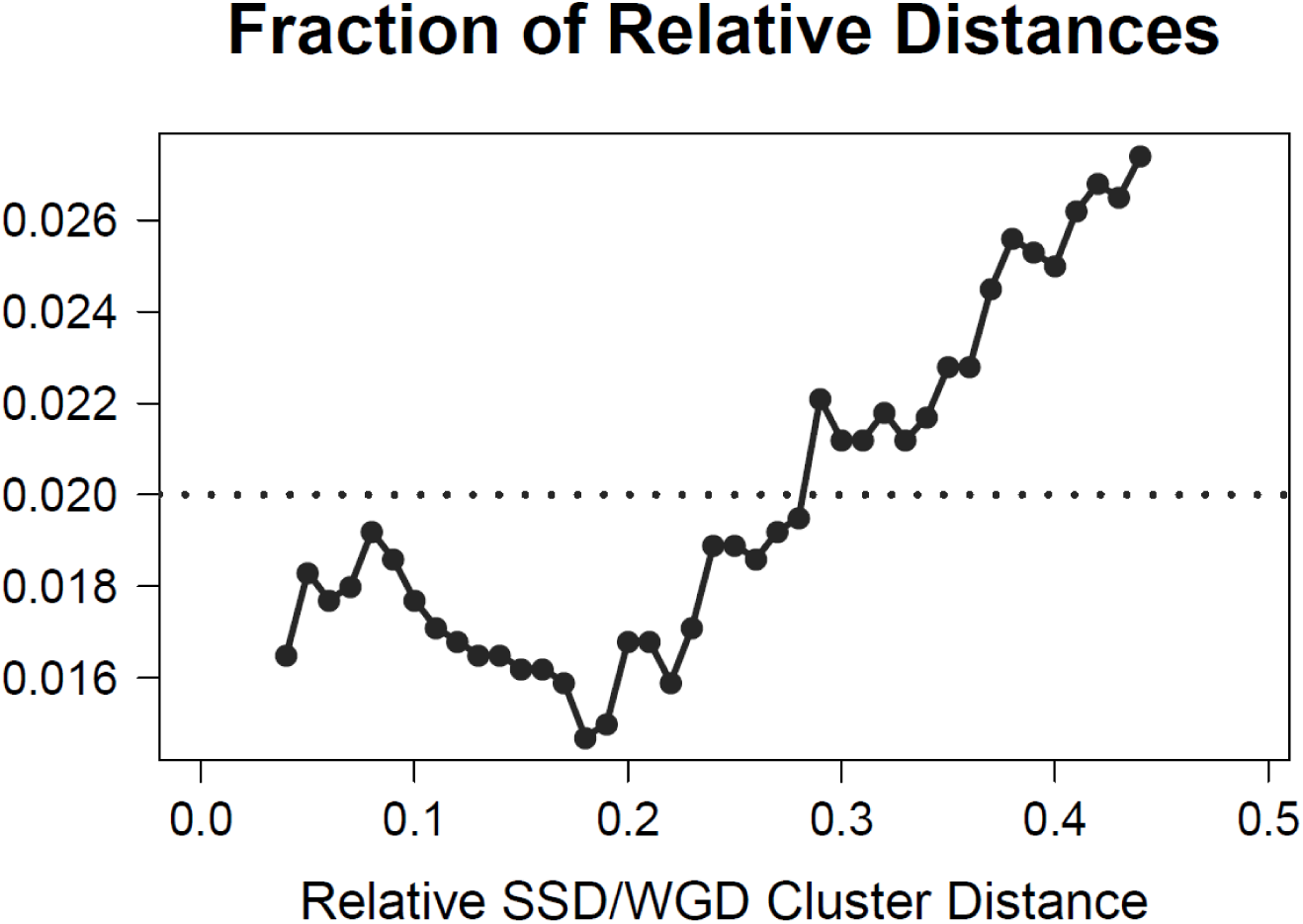
Fraction of relative distances between SSD-gene and WGD-gene clusters. Relative distance is defined as described in Favorov et al (2012) (see also Methods) with 0 corresponding to maximal co-occurrence and 0.5 to maximal distancing. Horizontal line corresponds to baseline fraction.

### Duplicate gene clusters occupy regions with different architectural preferences

The segregation of SSD and WGD, shown in Figure 1D, effectively divides the genome in three domains, one for each type of gene duplicate and the remaining complementary part of the genome, which is enriched for singleton, non-duplicated, genes. We went on to analyze particular spatial tendencies for these three compartments of the yeast genome. A first observation from Figure 1D suggests that SSD-clusters are enriched towards chromosomal edges, when compared to WGD-clusters and singletons. This is quantitatively supported by a comparison of the distributions of scaled distances from the chromosomal edges for genes residing in the three genomic domains (Supplementary Figure 1). Interestingly this tendency is also visible at the three dimensional level, with SSD clusters being preferentially positioned in the most peripheral regions of the genome according to the conformational model of (Duan et al. 2010) as opposed to more central positions for WGD ones (Supplementary Figure 2).

Duplicate genes are also found to have particular preferences in terms of size in both genic and associated non-genic sequences. Gene length is directly associated with functional complexity as longer genes may accomodate a larger number of functional domains (Lopes et al. 2021). In a similar way, the surrounding non-coding space may be associated with regulatory complexity, as longer promoters and gene upstream regions may provide the platform for a greater number of transcription factor binding sites (David et al. 2006). A simple comparison of gene lengths shows duplicate genes to be significantly longer than singletons (Figure 2A). This is partly explained by their enrichment in genes of greater age which are generally longer (Supplementary Figure 3) but also on the basis of the more complex evolutionary patterns of duplicate genes, which are more prone to maintain a large number of functions and thus their size (Vitkup et al. 2006; Costanzo et al. 2010; Kuzmin et al. 2020). Singleton genes have, on average, shorter lengths, independently of their position in the genome.

**Figure 2A.**
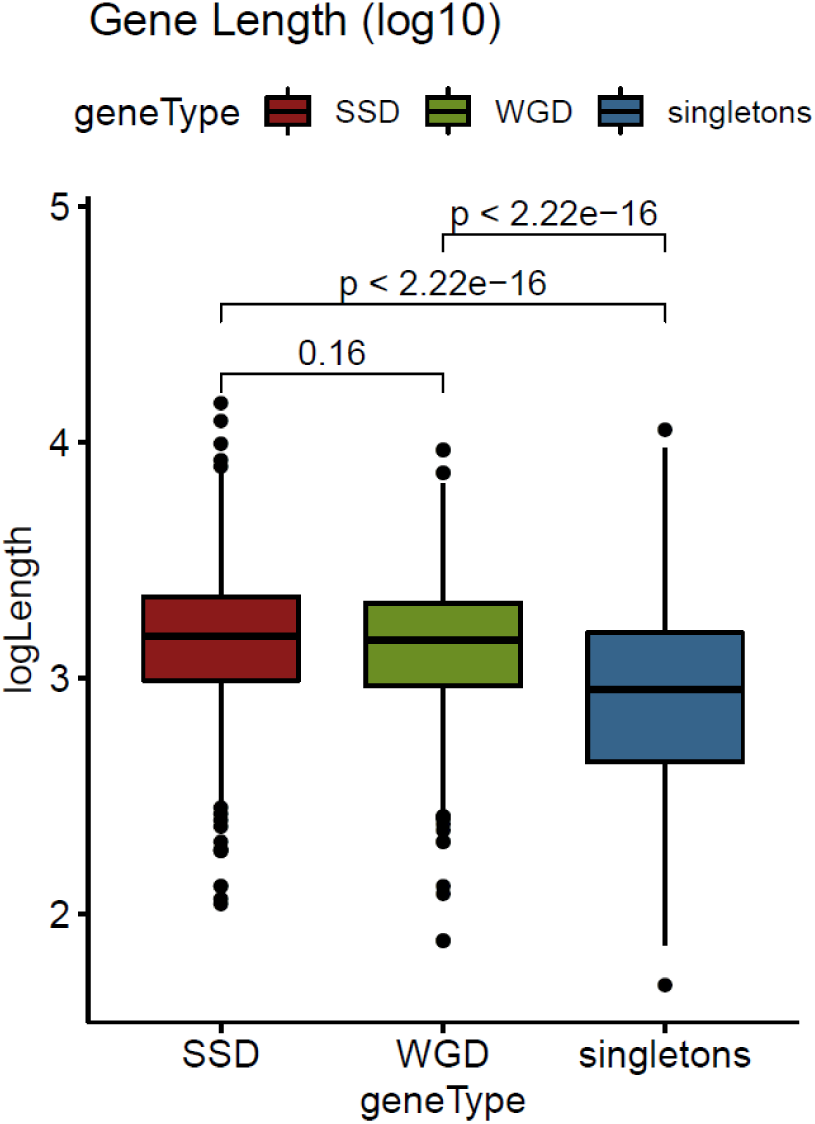
Distribution of gene lengths (log10-transformed) for SSD, WGD and singleton genes. Significance is denoted with p-values of a Mann Whitney test.

From an evolutionary viewpoint, the size of the proximal non-coding sequences may be equally important for functional divergence through the establishment of novel regulatory relationships. Adjacent genes that involve SSD duplicate pairs tend to have wider spacing (Byrnes et al. 2006) while duplicate genes with small distances are more likely to be deleted (Van Hoek and Hogeweg 2007). Both SSD and WGD are generally flanked by longer gene spacers (Supplementary Figure 4), but this is a feature that extends in duplicate gene clusters also affecting singletons.

We compared the coding sequence density (see Methods) for the clusters of SSD, WGD and singleton genes (Figure 2B) and found SSD clusters to be located in regions of the genome with the largest proportion of non-coding sequences. Moreover, both WGD and singleton genes were found to be embedded in areas of smaller coding density when found in the SSD-type gene clusters.

**Figure 2B.**
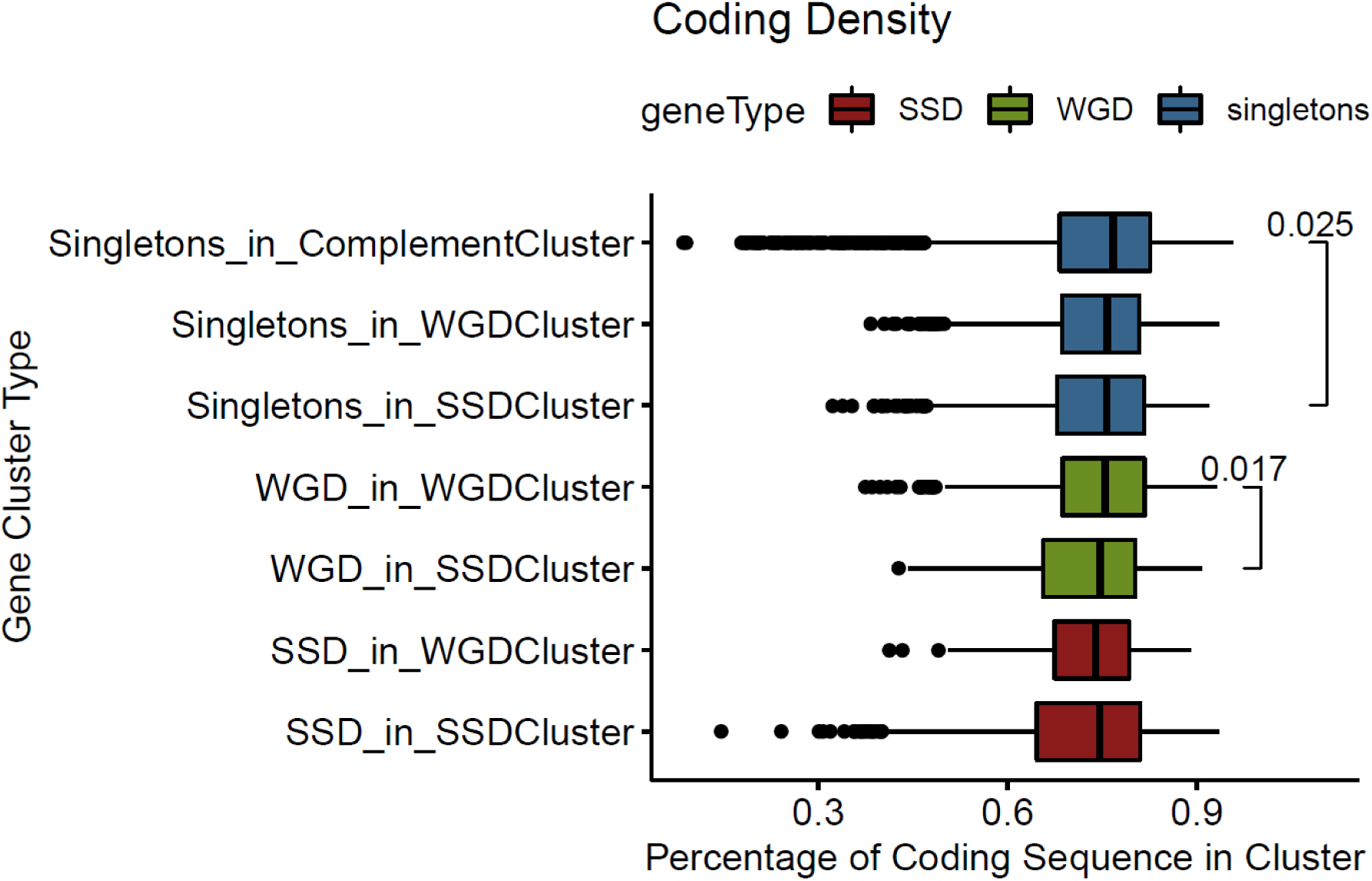
Distribution of coding density for regions around genes from different gene clusters (see text for details). Significant differences are denoted with p-values of a Mann Whitney test.

An explanation for this observation is the proximity of SSD gene clusters to the chromosomal edges. In the past, we have reported coding density to drop near the ends of chromosomes (Tsochatzidou et al. 2017) and indeed there is trend for longer gene spacers to be enriched towards the chromosomal edges (Figure 2C). These observations are suggestive of a general tendency of the genomic environment that doesn’t only affect genes of one particular category. In all, we find that the lengths of both coding and non-coding sequences, associated with gene duplicates, are dependent on the broader genomic area, in which these are embedded.

**Figure 2C.**
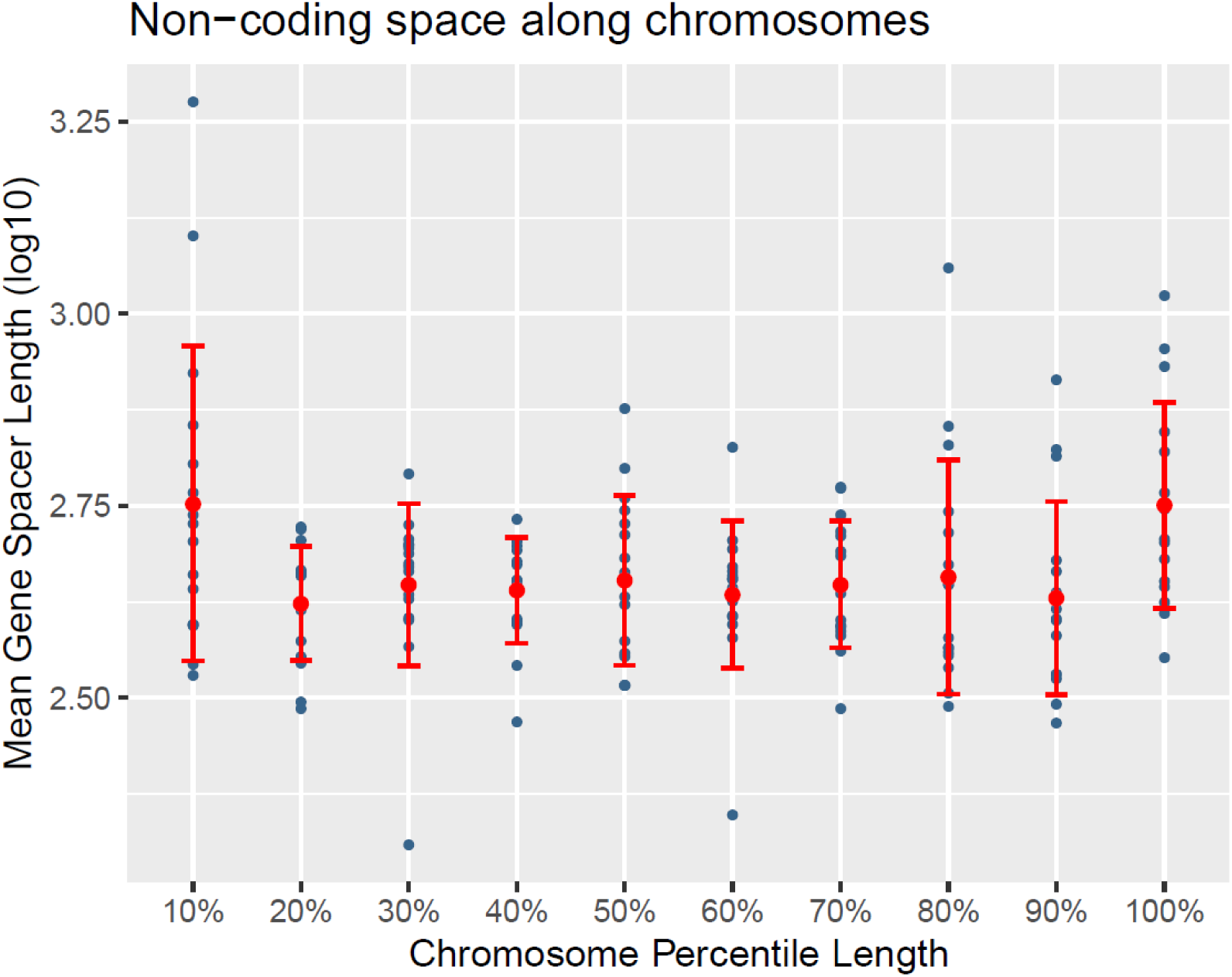
Mean gene spacer length (log10-transformed) along 10 quantiles in each of the sixteen yeast chromosomes. Dots correspond to mean values for the given percentile for each chromosome. Red bars correspond to standard deviation.

### SSD genes diverge at the regulatory level in a spatially-dependent manner

The increased size of non-coding spacers in gene duplicates and in particular SSD, prompted us to look closer into their gene upstream regions for possible sequence, structural and functional constraints. In order to assess sequence constraint, we created aggregate phastCons plots along the length of a region spanning 500bp either side of the transcription start site of each gene. Even though conservation is, in general, reasonably lower in the gene upstream region, we were surprised to find that the drop in conservation in the promoter region is sharper for both SSD and WGD genes, especially in the region immediately upstream of the gene’s transcription start site (Figure 3A).

**Figure 3A:**
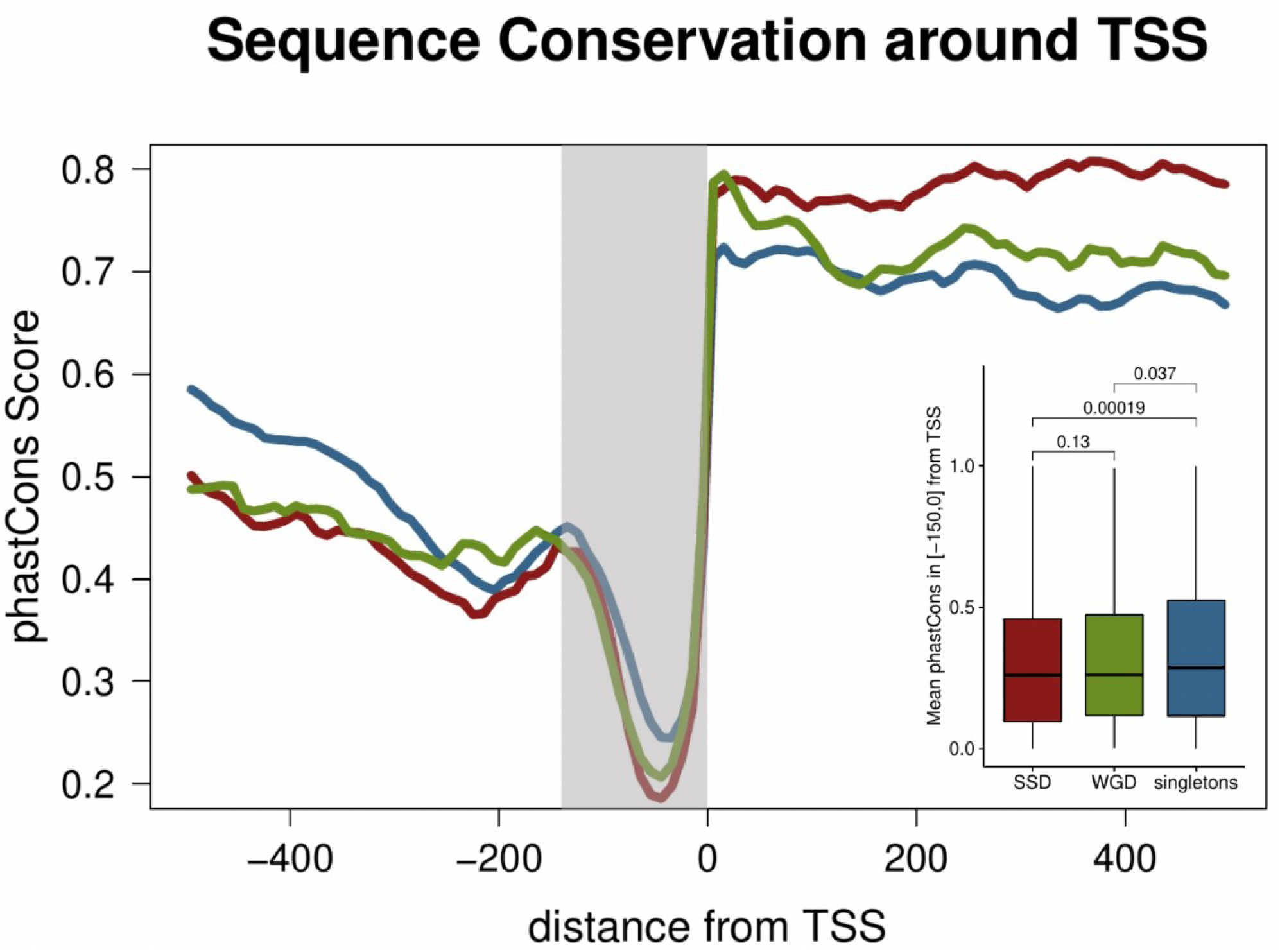
Aggregate mean conservation (phastCons) along a region spanning 500bps either side of TSS for singleton, SSD and WGD genes. Embedded boxplot represents mean phastCons scores for the highlighted, proximal promoter region (150bp upstream to TSS). Values in brackets correspond to p-values of a Mann-Whitney test.

This reduction in promoter sequence conservation, somewhat more pronounced in SSD compared to WGD, may be associated to more relaxed constraints in terms of gene regulation for duplicate genes against singletons, which is compatible with the general view of duplicate genes being more prone to diverge into new functions. What is more interesting, while coding sequence conservation appears to be independent from the region in which the gene is located, this is not the case for the reduced promoter conservation which follows a strong spatial pattern along the genome (Figure 3B).

**Figure 3B.**
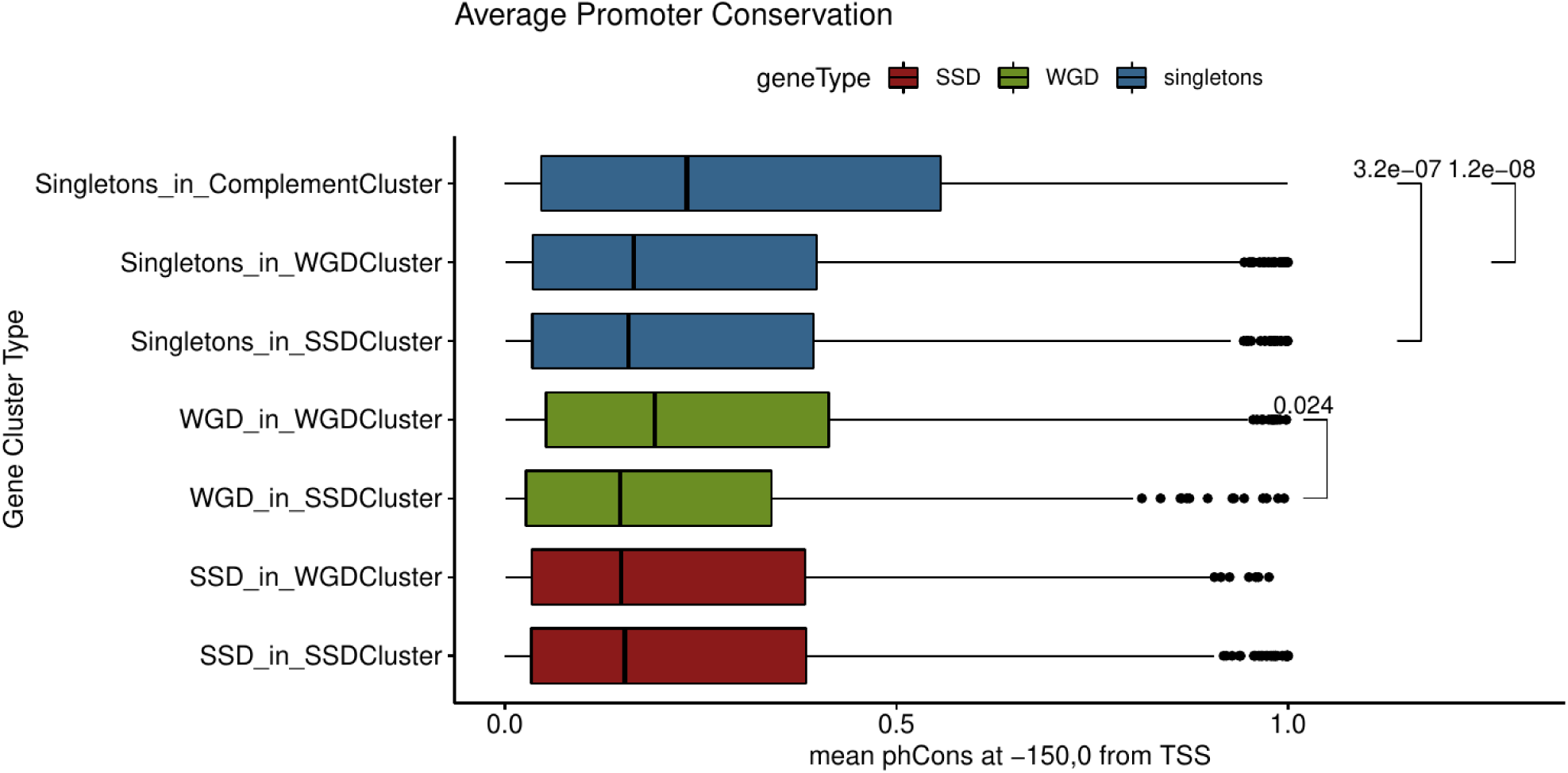
Distribution of mean conservation in the proximal promoter region (150bp upstream to TSS) measured as mean phastCons scores for different gene clusters. Values next to brackets denote p-values of a Mann Whitney test.

Singleton genes found in WGD clusters have less conserved promoters and this is further reduced in SSD clusters. It should be noted here that the estimated divergence between the duplicate gene pairs (Fares et al. 2013) is smaller in SSD genes compared to WGD ones, regardless of their position in the genome. Thus the apparent, more relaxed constraints in the promoter region are more likely to be associated with gene regulation and not overall higher divergence rates. We found support for this in the increased number of transcription factor binding sites (TFBS) identified at the promoters of different gene types. Both SSD and WGD have significantly greater numbers of conserved TFBS in an equally-sized gene upstream region (up to 300bp upstream of the TSS) when compared to singleton genes (Figure 3C).

**Figure 3C.**
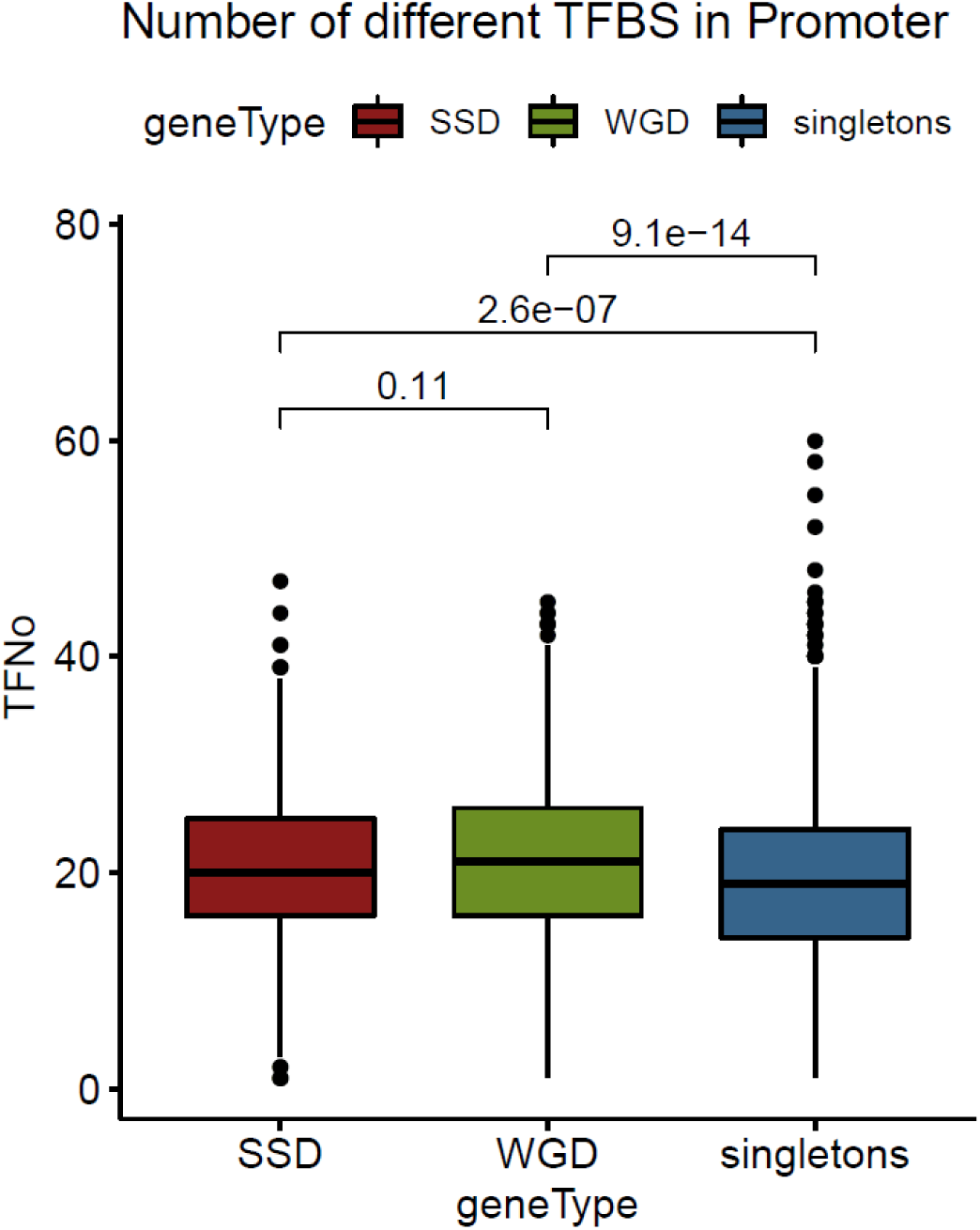
Distributions of numbers of predicted transcription factor binding sites (TFBS) in the gene upstream regions (300bp upstream to TSS) for SSD, WGD and singleton genes. TFBS obtained from a set of predictions based on comparative sequence analysis (MacIsaac et al. 2006).

Together, these observations suggest that, the combination of more extensive non-coding space in SSD clusters and relaxed sequence constraints in the promoters of the contained genes, may be the primary driving force for their functional divergence, occurring primarily at the regulatory level. Furthermore, there are strong indications of a spatially dependent relaxation of promoter constraints that occurs in areas enriched for SSD genes and which affects both singletons and WGD genes as well.

### Increased chromatin structural complexity for gene duplicates extends in broad genomic regions

We have recently described the organization of the yeast genome in extended regions with similar nucleosomal patterns which are, in addition, associated with a number of functional and regulatory characteristics (Nikolaou 2018). We used a public dataset (Jiang and Pugh 2009) to analyze the nucleosome occupancy patterns around the transcription start sites of SSD, WGD and singleton genes. The results showed marked differences in the gene upstream regions with both types of duplicate genes having more “shallow” nucleosome free regions (NFR), compared to a clear and deep NFR for singleton genes (Figure 4A).

**Figure 4A.**
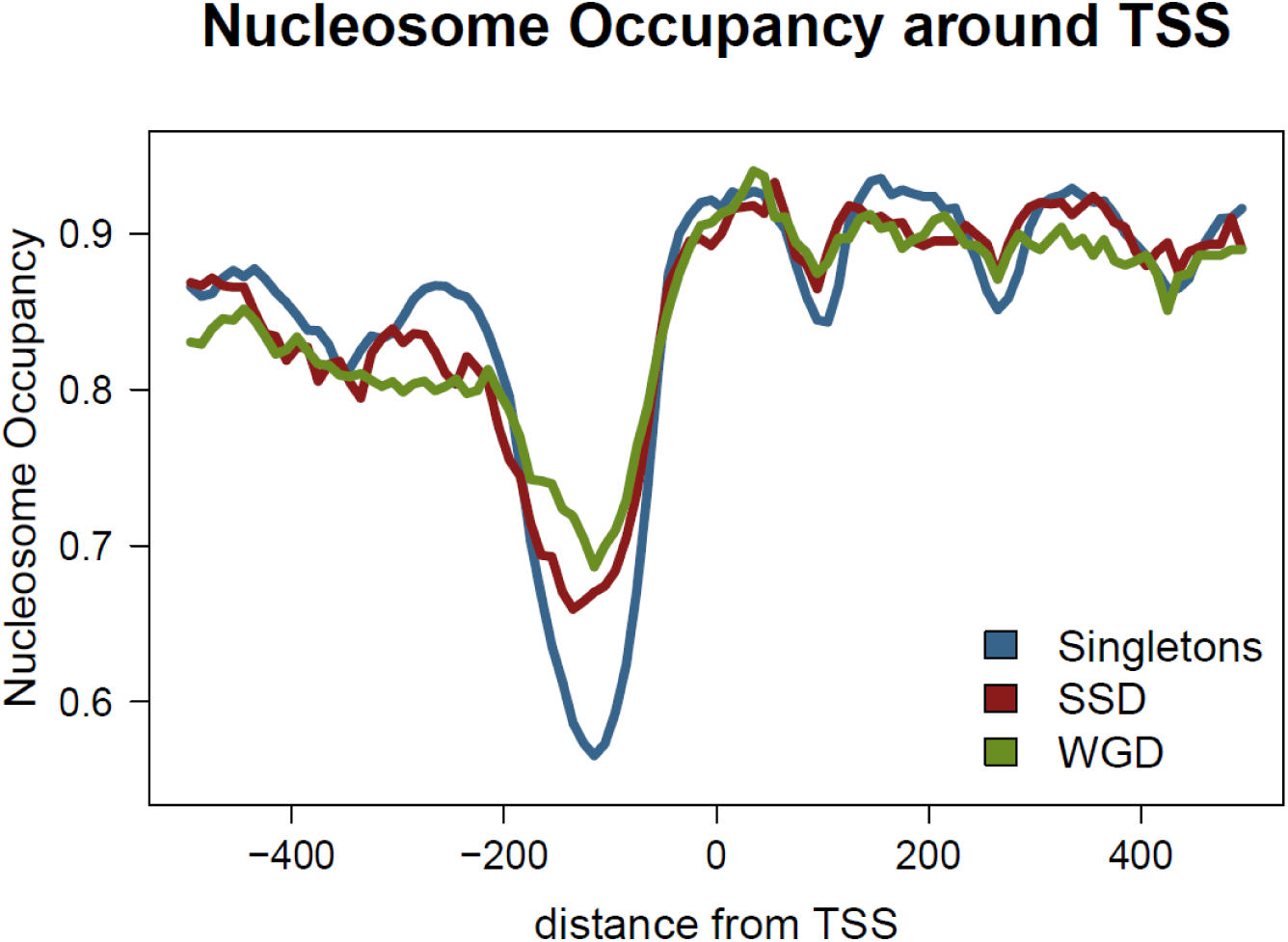
Mean nucleosome occupancy along a region spanning 500bps either side of TSS for singleton, SSD and WGD genes. Nucleosome occupancy was calculated in one hunded 10bp bins, as the fraction of the region overlapping a positioned nucleosome. Positions obtained from (Jiang and Pugh 2009).

Notably, this is not affected by the sample size, as the mean nucleosome occupancy at the promoter remains significantly lower even for a random selection of 1000 singleton genes (Supplementary Figure 5). Moreover, it appears to be position-independent as singleton genes have strong NFRs, regardless of the cluster they are found in.

Genes with strong NFR are generally subject to less complex regulation, as they do not require chromatin remodelling to allow for transcriptional activation by regulators (Yuan et al. 2005; Kaplan et al. 2009). They are thus enriched among constitutively expressed genes with more stable expression levels. Indeed, this is supported by our assessment of transcriptional level variability on the basis of data collected by the SPELL database (Hibbs et al. 2007), which show significantly smaller variability in mRNA levels for singleton genes compared to both SSD and WGD ones (Figure 4B). As the variability in mRNA levels is here measured between a broad spectrum of different conditions it may be seen as a proxy for expression plasticity, which is found to be increased in the case of duplicate genes. This comes in agreement with the observation of lower TFBS number in singletons.

**Figure 4B.**
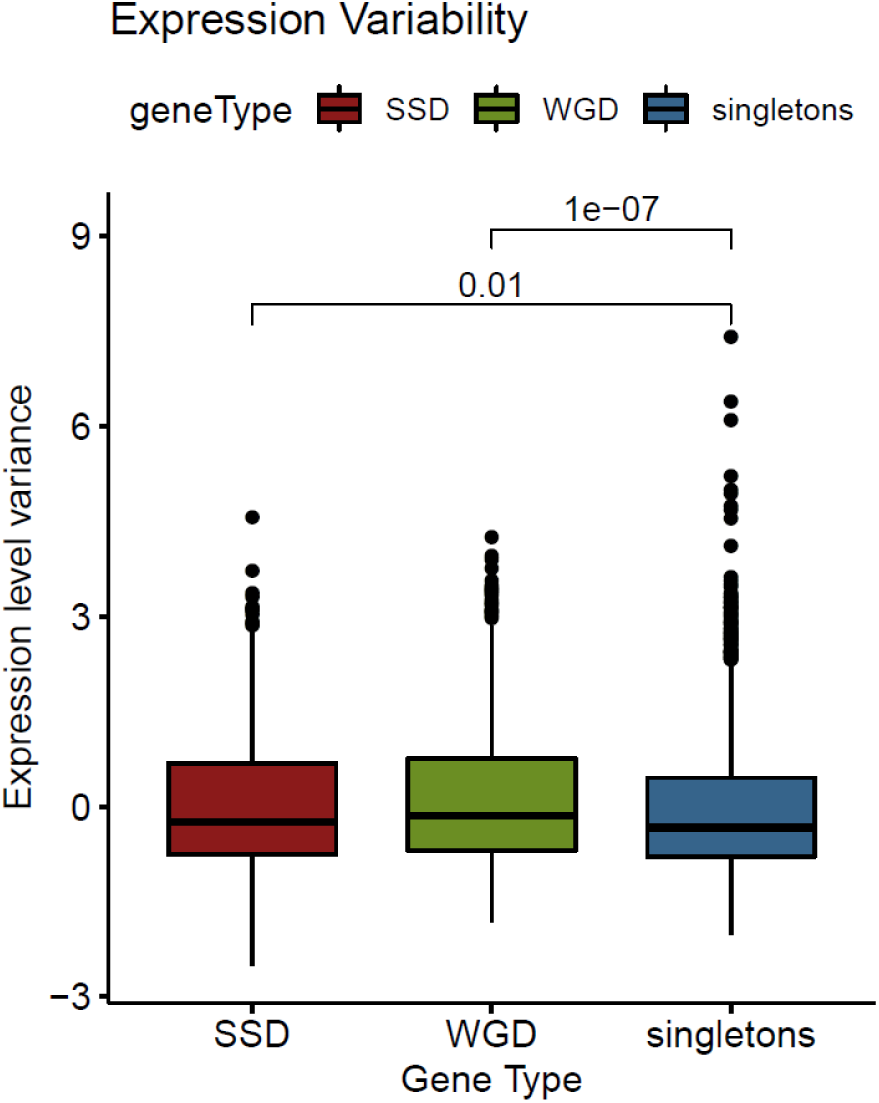
Distribution of expression variability for SSD, WGD and singleton genes. Expression variability was assessed as the z-score of the variance of gene expression values from the SPELL database. Values over brackets denote p-values of a Mann Whitney test.

The way sequence constraints may be affecting the positioning of nucleosomes has been the focal point of both experimental (Yuan et al. 2005; Segal et al. 2006) as well as computational works (Babbitt and Kim 2008; Kaplan et al. 2009). In the past we have suggested a simple method to assess structural constraints on the primary DNA sequence (Nikolaou et al. 2010) based on our own model for nucleosome positioning (Tilgner et al. 2009). A more recent work, using a deep learning model, has presented evidence of strong sequence constraints affecting nucleosome positioning in gene upstream regions (Routhier et al. 2020). We used both this deep learning model of mutation impact on nucleosome positioning, as well as our own simpler model of structural robustness (Nikolaou et al. 2010) (see Methods) to assess chromatin-related constraints around the TSS of duplicate genes and singletons. Both analyses produced similar patterns that show more elevated constraints in the gene upstream regions of duplicate genes compared to singletons. (Figure 4C, Supplementary Figure 6).

**Figure 4C.**
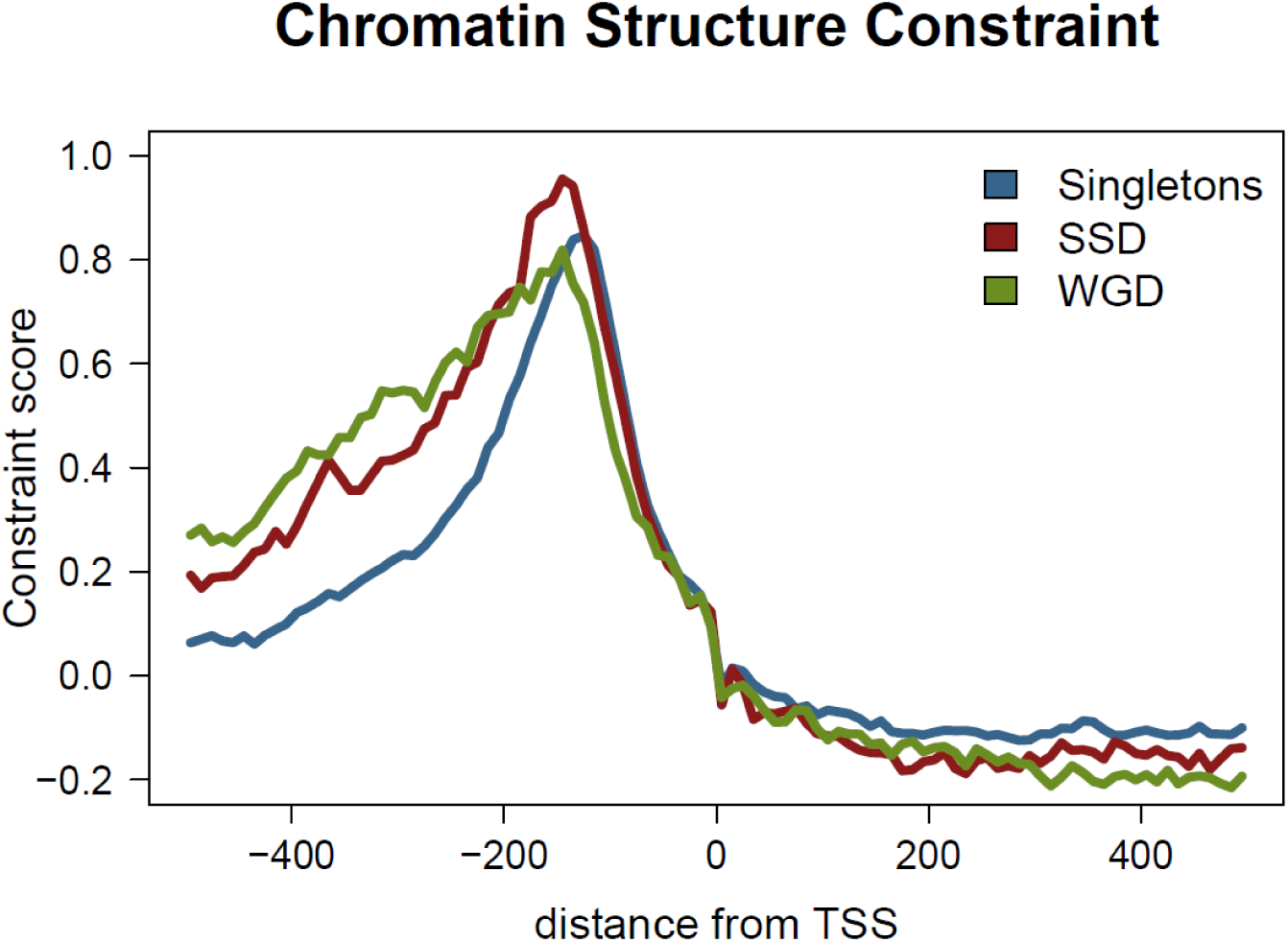
Chromatin structure constraint score as measured with the mutation score (Routhier et al. 2020) along a region spanning 500bps either side of TSS for singleton, SSD and WGD genes.

In the case of duplicated genes the constraints were also more extended upstream, which may be related to the overall larger size of their gene upstream regions (see above). The increased structural constraints of gene duplicates are also spatially associated, in a way similar to sequence constraints. What should be noted is that most of the constraint observed for singletons is due to singleton genes found in SSD and WGD clusters (Supplementary Figure 7), as the mean structural conservation of singletons in SSD/WGD clusters is significantly higher than the one of singletons in the rest of the genome (Supplementary Figure 8).

Together, these observations point towards strong structural constraints in the promoters of genes that are found in the areas of the genome, where duplicate genes are preferentially positioned.

### Structural constraints at the promoter are independent of sequence conservation but may be shaping the gene expression of gene duplicates

The existence of structural constraints in duplicate gene clusters appears to contrast the relaxed sequence conservation in their promoters, which we show earlier. We went on to examine the association between the two and we were not surprised to observe a significant negative correlation (Spearman’s rho = -0.229, p<=10^-57^). This inverse relationship between sequence conservation and structural constraint at the gene promoters appears to be a general characteristic (Figure 5A) and may indicate that relaxation at the sequence level is counteracted at the level of structure.

**Figure 5A.**
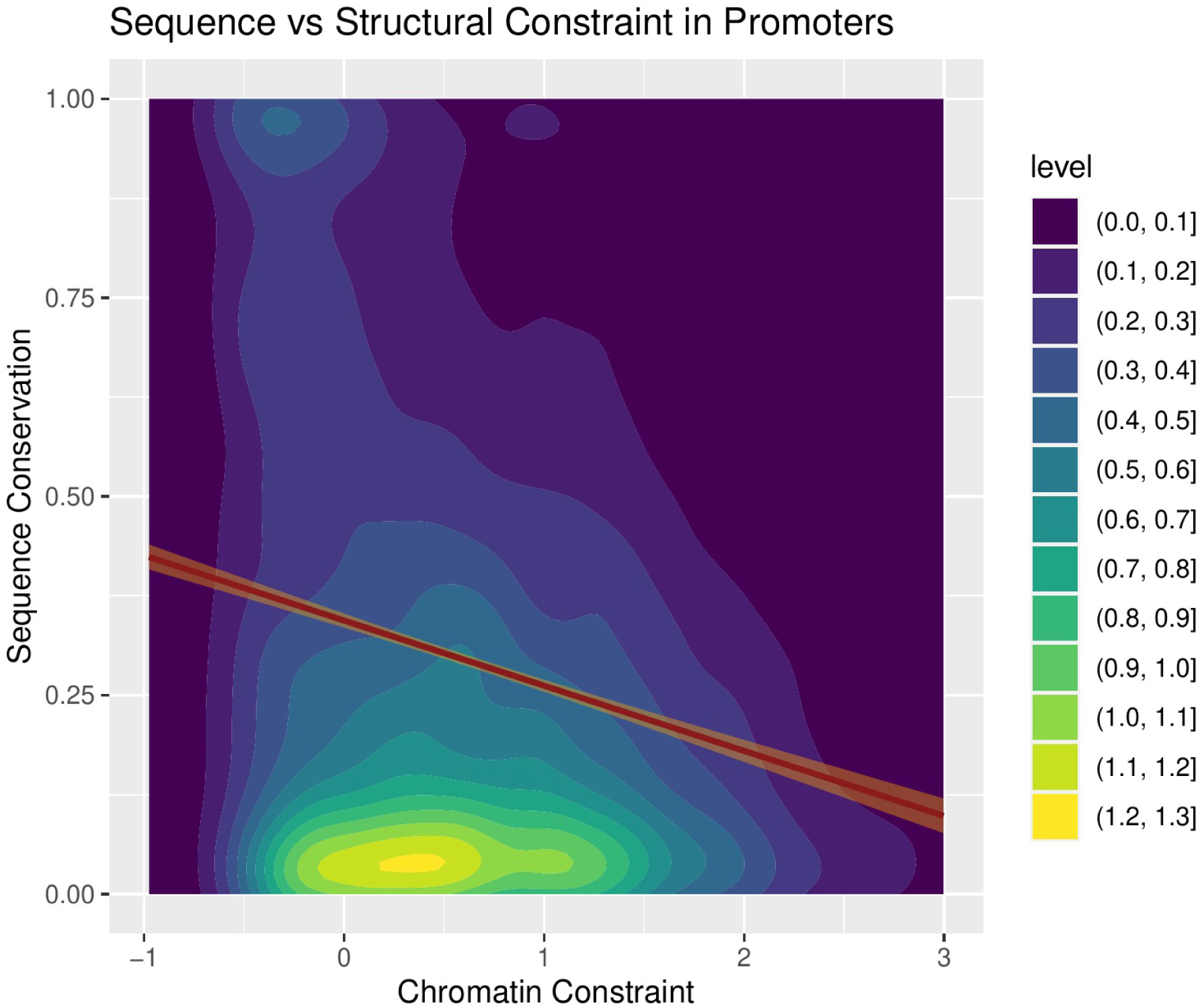
Two-dimensional density plot of sequence constraint (as mean conservation score) versus structural constraint (as mean mutation score by (Routhier et al. 2020)) for the same regions of 200bp to 50bp upstream of the TSS for all yeast genes (Spearman’s rho = -0.229, p<=10^-57^). Level values correspond to density of genes in the two-dimensional space. Red line corresponds to a linear regression fit, with orange band corresponding to standard error.

What is more, it denotes two distinct promoter architectures. On one hand there are more conserved, small promoters with clear nucleosome-free regions pertaining to singleton genes, which overall have less complex regulation and smaller expression variability. On the other, broader promoters with stronger structural constraints and more complex nucleosomal patterns are representative of duplicate genes, which, in turn, have more complex regulatory patterns and greater transcriptional plasticity.

The co-regulation of gene duplicates has been shown to be dependent on both their linear (Lan and Pritchard 2016) and three-dimensional proximity (Ibn-Salem et al. 2017) but the role of chromatin structure in the modulation of gene regulation has not been investigated. We analyzed the effect of nucleosome positioning on gene duplicates by comparing the structural similarity of the sequence around the TSS with an adjusted co-expression score for each duplicate gene pair. Structural similarity was calculated as the Pearson correlation coefficient of nucleosome positioning patterns for an area of 1000bp symmetrically flanking the TSS (see Methods). The adjusted co-expression score (ACS) was obtained from the SPELL database (Hibbs et al. 2007) as a weighted correlation of gene expression levels estimated for >2400 different experimental conditions. We found a small, yet significant correlation for both types of gene duplicates (all duplicates Spearman’s rho = 0.089, p=0.0035), which was even stronger for SSD (SSD Spearman’s rho = 0.109, p=0.015) (Figure 5B).

**Figure 5B.**
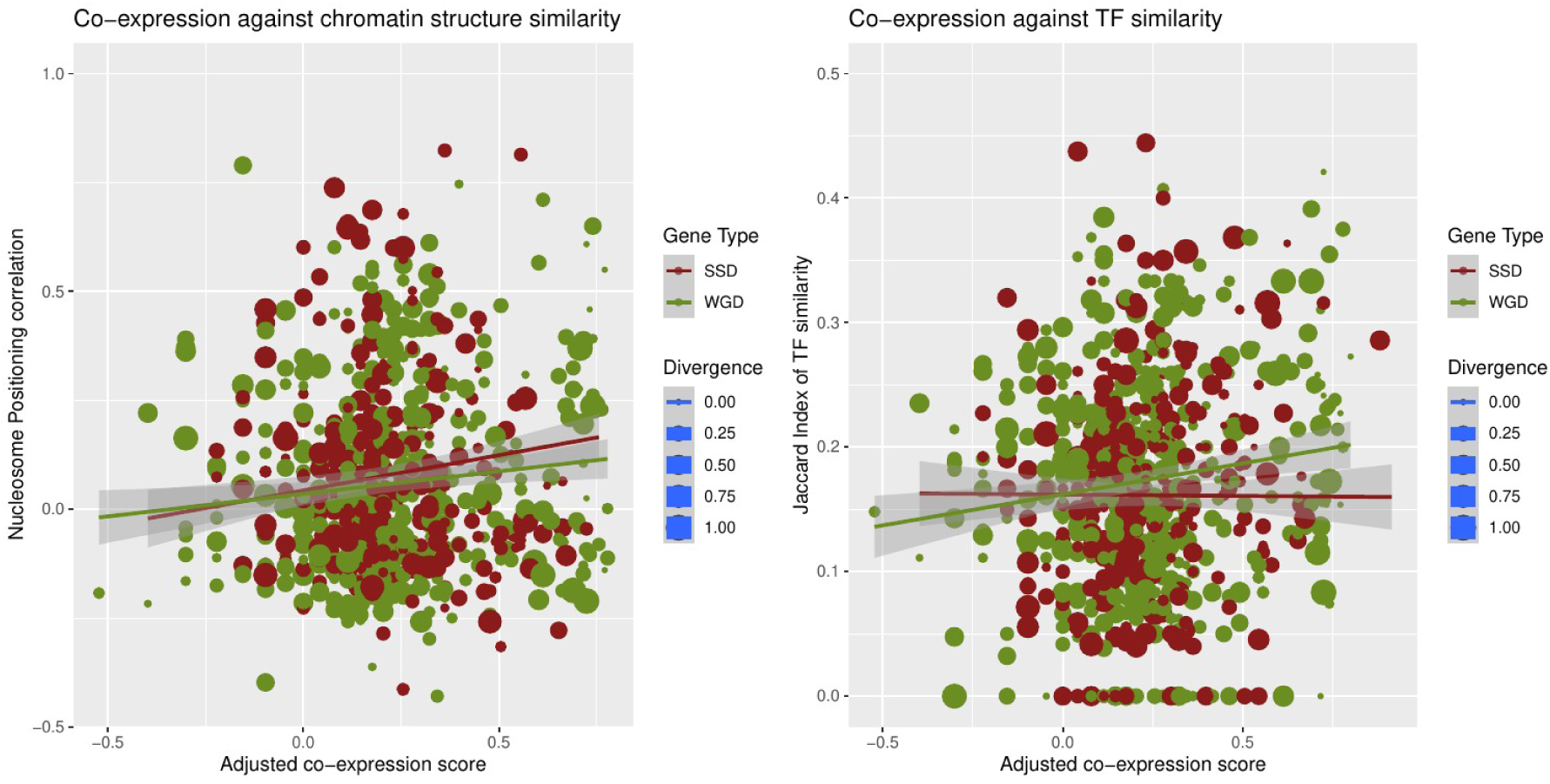
Left: Two-dimensional scatterplot of gene pair co-expression, measured as scaled Adjusted co-expression Score (obtained from SPELL) against nucleosome positioning correlation, measured as the Pearson correlation coefficient of nucleosome occupancy profiles between gene pairs. Right: Gene pair co-expression against TF similarity measured as the Jaccard index of TFBS found in the promoters of each gene pair. Red: SSD gene pairs, Green: WGD gene pairs. Size of points proportional to the estimated diversity between the genes in each pair.

This correlation between structural similarity and gene co-expression was independent of the sequence divergence between the gene pair. A number of studies have suggested that divergence is more pronounced at the level of regulation than at that of gene sequence or function (Byrnes et al. 2006; Wapinski et al. 2007). In order to assess this association, we performed the same analysis comparing the adjusted co-expression score with the regulatory similarity, as assessed with the Jaccard similarity of common TFs having a conserved binding site at the genes’ promoters (see Methods). The effect of regulatory similarity on co-expression is comparable (all duplicates Spearman’s rho = 0.096, p=0.0017) but in this case it is the WGD genes that show the stronger association (WGD Spearman’s rho = 0.119, p=0.0053) (Figure 5B). Together, these results suggest that the local chromatin environment may shape the expression patterns of duplicate genes in a way that is as strong as the one exhibited by transcription factor binding. Moreover, they paint a complex picture of the modulation of functional divergence. SSD genes appear to be more dependent on structural properties than TF binding, while the opposite is the case for WGD ones (Figure 5B).

### Singleton gene functional and regulatory properties are dependent on their location in the genome

Combined, our observations suggest that a set of spatial, structural and regulatory properties define different genomic “niches” that are occupied preferentially by SSD and WGD duplicates. Given that a number of these properties affect the broader environment and are shared by proximal singleton genes, one hypothesis is that singletons found in duplicate gene clusters may partly share the evolutionary history of nearby duplicate genes. This may be particularly interesting for the case of singletons in SSD gene clusters, as SSD genes occupy genomic regions which appear to be more prone to complex regulation (longer non-coding regions, stronger chromatin constraints and more relaxed sequence promoter conservation). We wanted to examine the possibility that a set of singleton genes, sharing SSD-like properties, may constitute remnants of a duplication event and which may be, in part, maintaining some characteristics that distinguish them from genes that have not undergone gene duplication, at least not recently.

One way to examine this is by analyzing the similarity of genomic sequences that contain some residual similarity with existing genes and may thus represent gene “relics”, products of a recent duplication event, which have acquired a sufficient amount of substitutions to render them indistinguishable from intergenic DNA. We used a dataset of 124 such relics from the yeast genome, bearing similarity with 149 distinct genes, identified through a stringent sequence similarity analysis (Lafontaine et al. 2004). Genes with similarity to gene relics were preferentially positioned in SSD gene clusters. In addition, they were enriched in both SSD and singletons, but not WGD genes residing in these clusters (Figure 6A). This is suggestive of more frequent duplication events in the areas of the genome lying towards the chromosomal edges, with longer intergenic spacers. It also indicates that a significant proportion of singleton genes found in these regions constitute remnants of small-scale duplications. This hypothesis is further supported by the fact that the relics themselves are preferentially found in SSD gene clusters, with an observed/expected ratio, o/e=1.69 (p=0.02), compared to a clear depletion in WGD clusters (o/e=0.45, p=0.04) (Supplementary Figure 9).

**Figure 6A.**
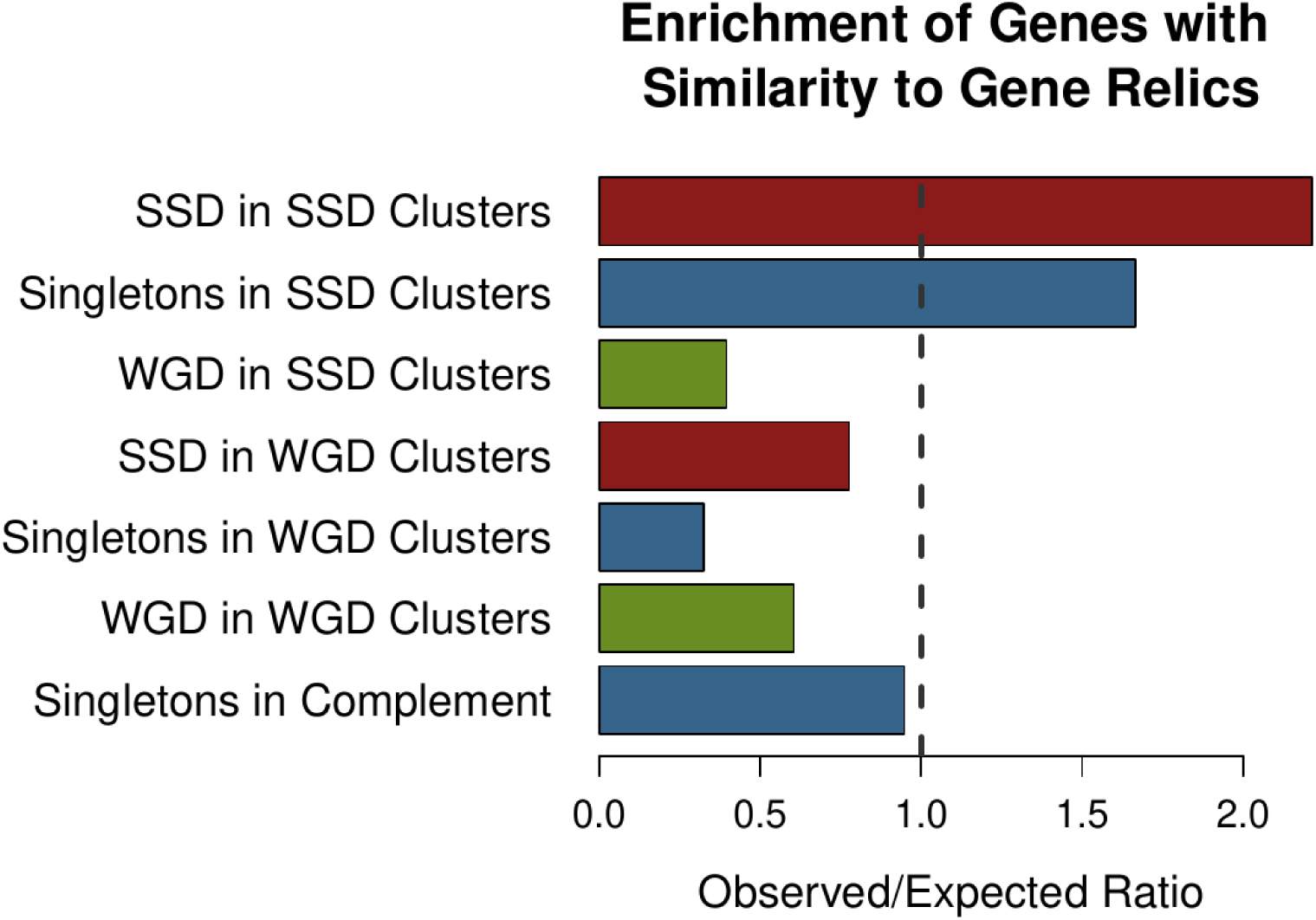
Relative enrichment of genes with similarity to gene relics in the different cluster categories

Reflections of the evolutionary history of such “vestigial” singleton genes may be found in their functional properties. According to the definition of functional entanglement by (Kuzmin et al. 2020), functionalities associated to specific protein domains may be structurally constrained and thus restrict the way genes evolve. Entangled, constrained functions do not allow for the evolutionary divergence and sub-functionalization and a gene duplicate pair with highly constrained functions is more likely to revert to a singleton state. We used the predicted PFAM domains as a proxy for protein functional domains and assessed the percentage of the gene covered by a known PFAM as a measure of its functional complexity. We found that singletons in WGD and (even more) in SSD gene clusters have significantly increased fractions of their length assigned to a functional domain (Figure 6B) even though they are of similar length (compare with Supplementary Figure 2).

**Figure 6B.**
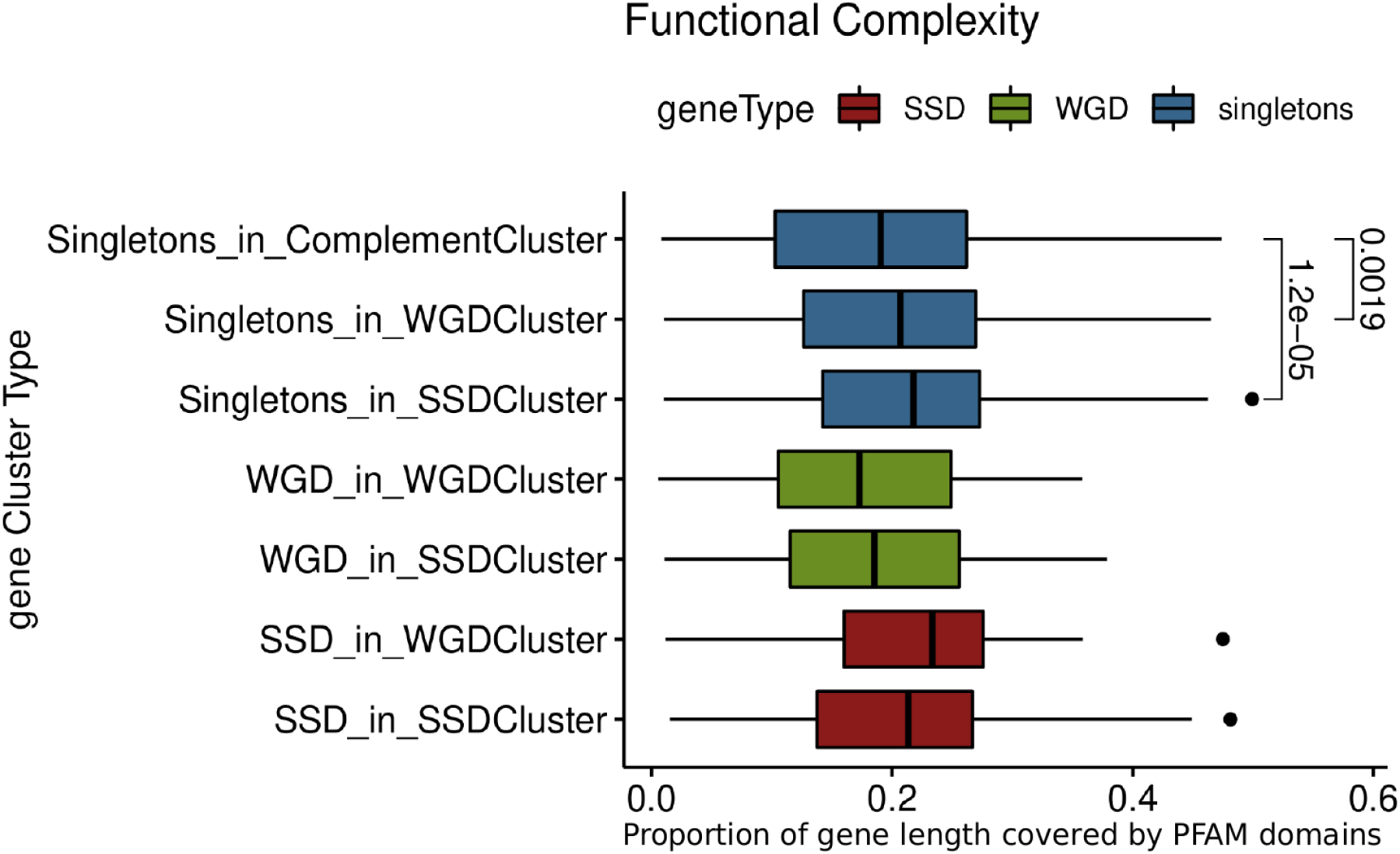
Distribution of functional complexity measured as the proportion of gene length attributed to a known PFAM domain for genes from different gene clusters. Values next to brackets denote p-values of a Mann Whitney test.

Increased functional complexity for singletons from gene duplicate clusters may suggest greater overall involvement in protein-protein interactions (PPI). As already suggested by (Fares et al. 2013), gene duplicates tend to have more PPIs. This may be attributed to a number of characteristics that we have identified above such as increased protein sequence length, regulatory complexity, expression variability and functional entanglement. It was thus interesting to see that singleton genes residing in duplicate clusters also tend to have increased numbers of PPIs (Figure 6C).

**Figure 6C.**
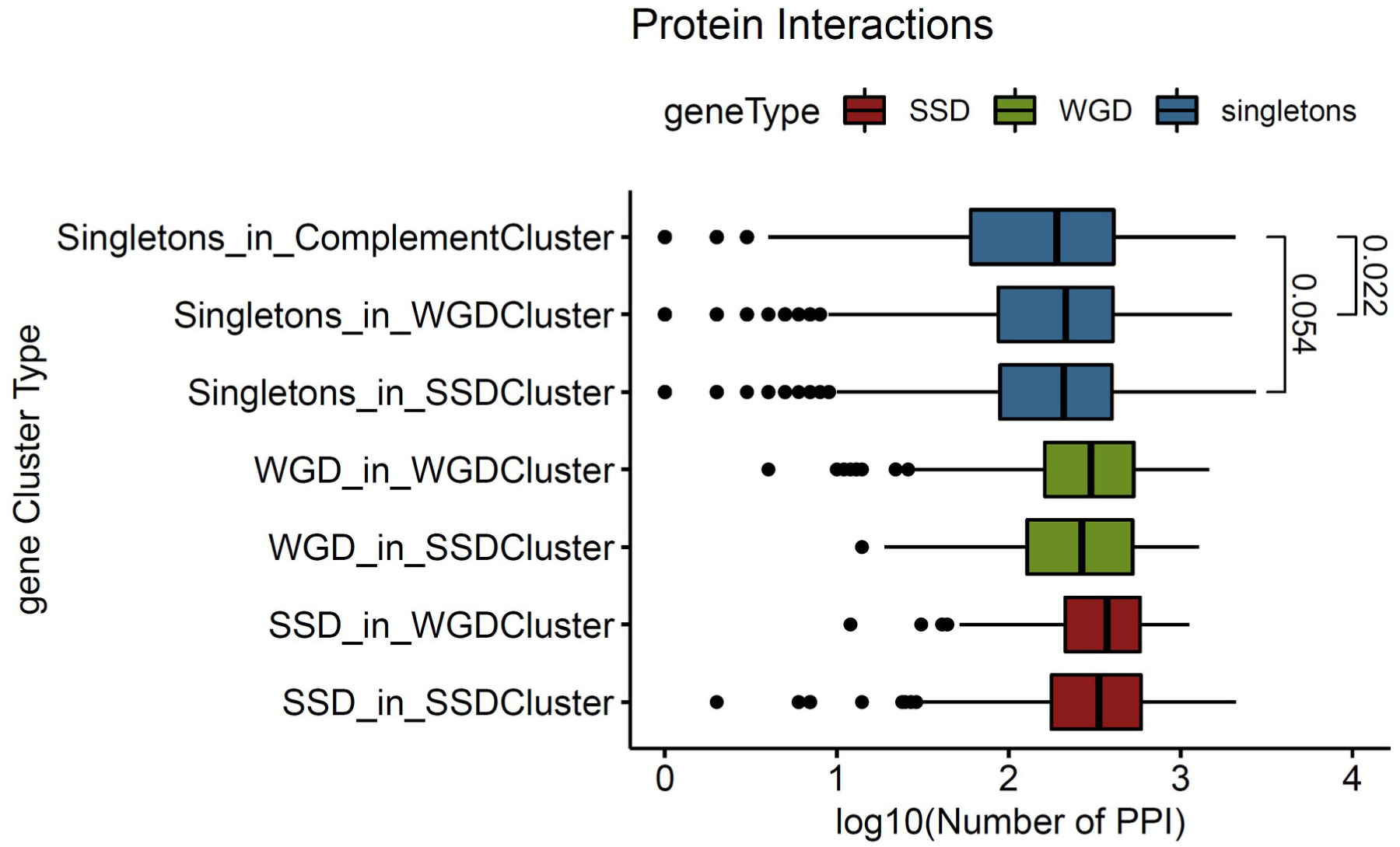
Distribution of the number of protein-protein interactions (log10-transformed) for genes from different gene clusters. Values next to brackets denote p-values of a Mann Whitney test.

Interaction preferences are not limited to protein-protein but extend to regulatory relationships. We calculated the relative enrichments of transcriptional regulator binding sites in genes belonging to each category and found the profiles of singletons in SSD clusters bearing stronger similarities with SSD genes than with singletons overall (Supplementary Figure 10), being enriched in binding sites of XBP1, MET4 and STB4 among others.

As enrichments values can also be biased towards over-represented regulators in the dataset, we further explored regulatory interactions through an association rules analysis, that aims to capture significant associations by better controlling for very common regulators (see Methods). By this point, we were not surprised to see that the association rules learned were also position-specific, with singleton and SSD genes exhibiting regulatory associations largely defined by the region in which they were located (Figure 6D). A number of regulators, including PDR3, ARR1 and ZAP1 were found to be strongly associated with both singletons and SSD genes when found in SSD clusters, suggesting the existence of pervasive, position-specific regulatory preferences.

**Figure 6D.**
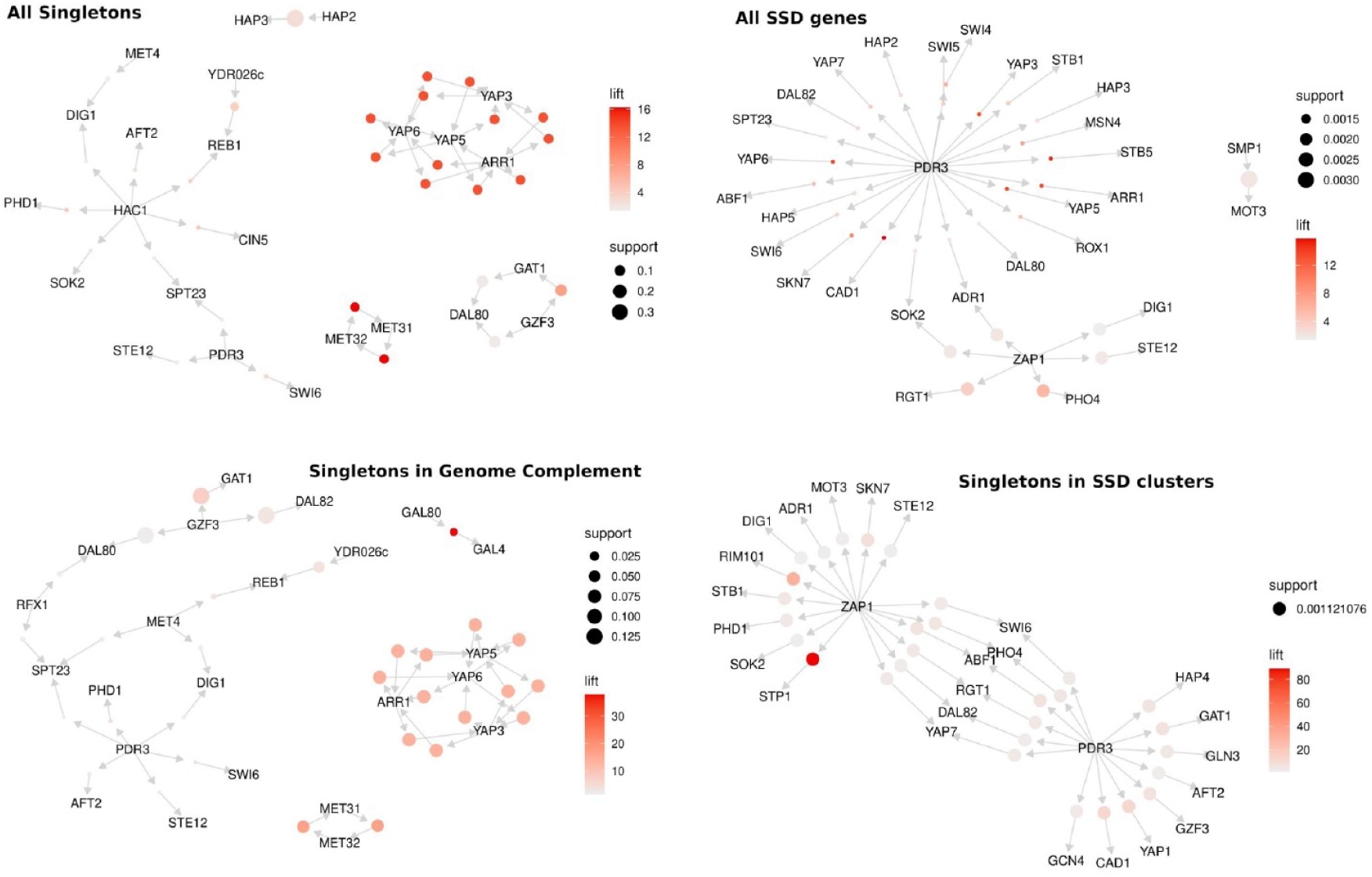
Association rule networks created for distinct sets of genes against transcriptional regulators whose binding sites were found in the genes’ promoters. Networks describe the top 10% most significant associations between regulators that share gene targets, omitting these targets for simplicity.

## Discussion

Recent advances at both experimental and theoretical levels have provided evidence for the structural organization of eukaryote genomes (Dekker et al. 2013; Tanay and Cavalli 2013; van Steensel and Furlong 2019). Even though, yeast has a rather small and dense genome in comparison to multicellular eukaryotes, a certain degree of organization exists at both one (Janga et al. 2008; Henikoff et al. 2011; Kouzine et al. 2013) and three dimensions (Duan et al. 2012; Rutledge et al. 2015). The existence of an underlying conformational genome scaffold alludes to the possibility of a concomitant functional compartmentalization. Aspects of such a compartmentalized view of the yeast genome may be seen at the localized chromatin structure of the genes’ promoters (Nikolaou 2018) as well as at more generalized “architectural” properties such as gene spacing and promoter complexity (Tsochatzidou et al. 2017).

In this work, we show that a certain degree of functional segmentation in the yeast genome is strongly associated with the localized potential for genomic innovation. We find gene duplicates to be concentrated in specific areas of the genome and, moreover, that they strongly segregate depending on their mode of duplication. Small-scale duplicates, which are in general more prone to diverge into acquiring novel functions, are preferentially located towards the chromosomal edges. This is consistent with the view of small-scale duplicates being intrinsically faster evolving, which has been reported for primates (O’Toole et al. 2018). Whole-genome duplicates, that mainly subfunctionalize, are positioned in more central genomic regions. This positional segregation offers likely explanations for the evolutionary fate of both gene duplicates as well as the singleton genes that are found in the same areas.

A sufficient amount of genome space is required for a duplicate to first be created without interrupting nearby genes and regulatory elements. In addition, it is more likely to be maintained if it is long enough to accommodate a number of functions that will allow its divergence (Kuzmin et al. 2020). Both of these conditions are met in the regions of the chromosomal edges which are less gene dense, with long non-coding spacers, which thus constitute a genomic “niche” that is more permissive for genomic innovation. Reflections of this may be seen in the functional enrichment of genes found in SSD clusters, with a strong over-representation of transmembrane proteins and transporters. Genes found in WGD clusters are, in contrast, primarily enriched in basal metabolic pathways and functions (Supplementary Figure 11). A general association of SSD genes with more specialized functions associated with stress response and specific conditions is also supported by the transcriptional regulator enrichments (Figure 6D, Supplementary Figure 10), with the majority being related to stress response and the use of alternative energy and nutrient resources. A plausible scenario for the confinement of such genes into specific chromosomal areas is that it may be confering an advantageous genomic “division of labour”, whereby novel functions may be explored in certain parts of the genome, minimizing the possibility of interference with regions hosting more stably expressed, constitutive genes. Stress response genes are known to be more likely to revert to singleton state when duplicated (Wapinski et al. 2007). In this sense SSD gene clusters, located at the chromosomal edges, would correspond to dynamic regions of high duplicate turnover, driving genomic innovation. An additional, strong indication for this is the enrichment of gene “relics” in these areas.

The mechanism, through which the exploration of genomic innovation takes place is far from understood, but our findings point to chromatin being an unexpected, yet crucial property in this respect. Both SSD and WGD genes have singular nucleosomal architectures at their promoters, that are both quantitatively and qualitatively different from the ones of singleton genes, suggesting that a structurally complex promoter is a primal property of gene duplicates. This is reflected on the increased and more extended structural constraints observed not only for gene duplicates, but also, residually, for singletons found in duplicate clusters. This finding appears at first counter-intuitive, especially when one sees its inverse correlation with gene conservation at the promoters (Figure 5A). However, we should consider that the structural constraints discussed herein are related to the maintenance of nucleosome positioning, which is only loosely associated with specific sequence signatures (Kaplan et al. 2009; Nikolaou et al. 2010; Ioshikhes et al. 2011). This means that a gene promoter’s structural profile may be modulated with some minimal prerequisites of sequence constraint, in a way that is permissive for the exploratory process of genomic innovation. Expression and eventually functional divergence may thus be achieved with a more complex promoter structure with minimal changes at the level of gene sequence, at least at the early stages of the divergence process. This is supported by a number of observations including the higher correlation of gene co-expression with nucleosome positioning similarity than with sequence divergence (Figure 5B). Interestingly, the correlation between gene co-expression and nucleosome positioning is almost 3-fold increased for duplicate genes lying close to the chromosomal edges compared to those being more central (cc=0.198 for duplicates lying within 10% of the length of a chromosomal arm from the edge as opposed to cc=0.068 for genes lying at the opposite 90%). Our findings, related to the nucleosome positioning structure, structural constraints and their relationship with expression divergence are highly indicative of local chromatin structure being a “soft” constraint, which acts permissively for the exploration of novel functionalities, without compromising the duplication and reverting into singleton. Such events are obviously the majority even in the “permissive” SSD niches as indicated by the similarity of singletons in these areas with gene relics (Figure 6A).

Overall, our observations regarding the properties of singleton genes that co-localize with gene duplicates, are supportive of our main hypothesis, that specific genomic niches are more tolerant to gene duplicates. New duplication events are therefore preferentially taking place in genomic subcompartments with specific properties. Singletons found in these subcompartments residually carry many of the attributes of gene duplicates and could be speculated to constitute remnants of recent duplication events. In all, our findings point towards an architectural segregation of function, regulation and evolvability in the yeast genome, which, buffered by chromatin structure, creates permissive environments for both neo- and sub-functionalization and thus creates preferential “niches” for gene duplicates.

---

**Supplementary Figure 1.**
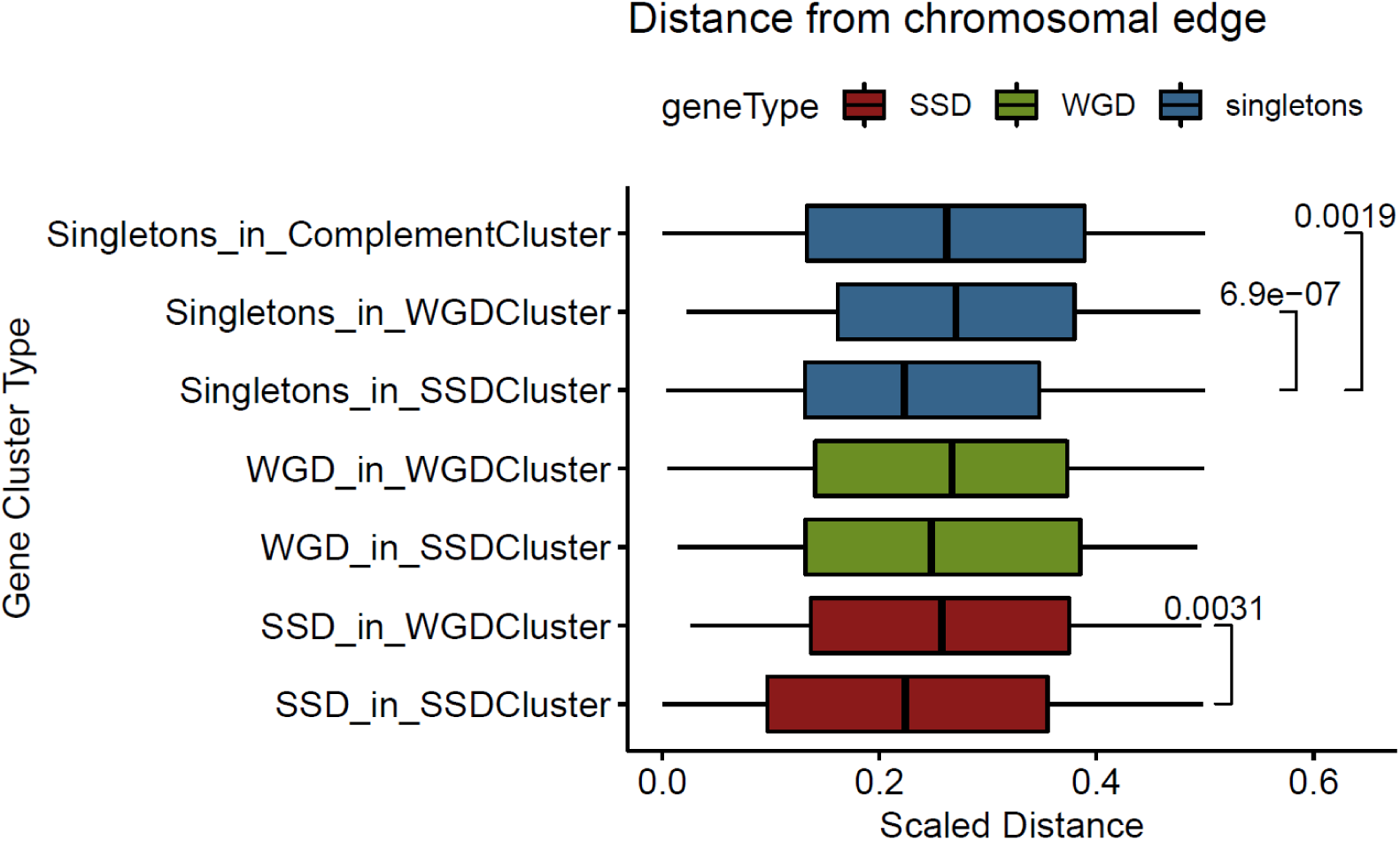
Distribution of scaled distances of gene clusters from the most proximal chromosomal edge. Significant differences are denoted with p-values of a Mann Whitney test.

**Supplementary Figure 2.**
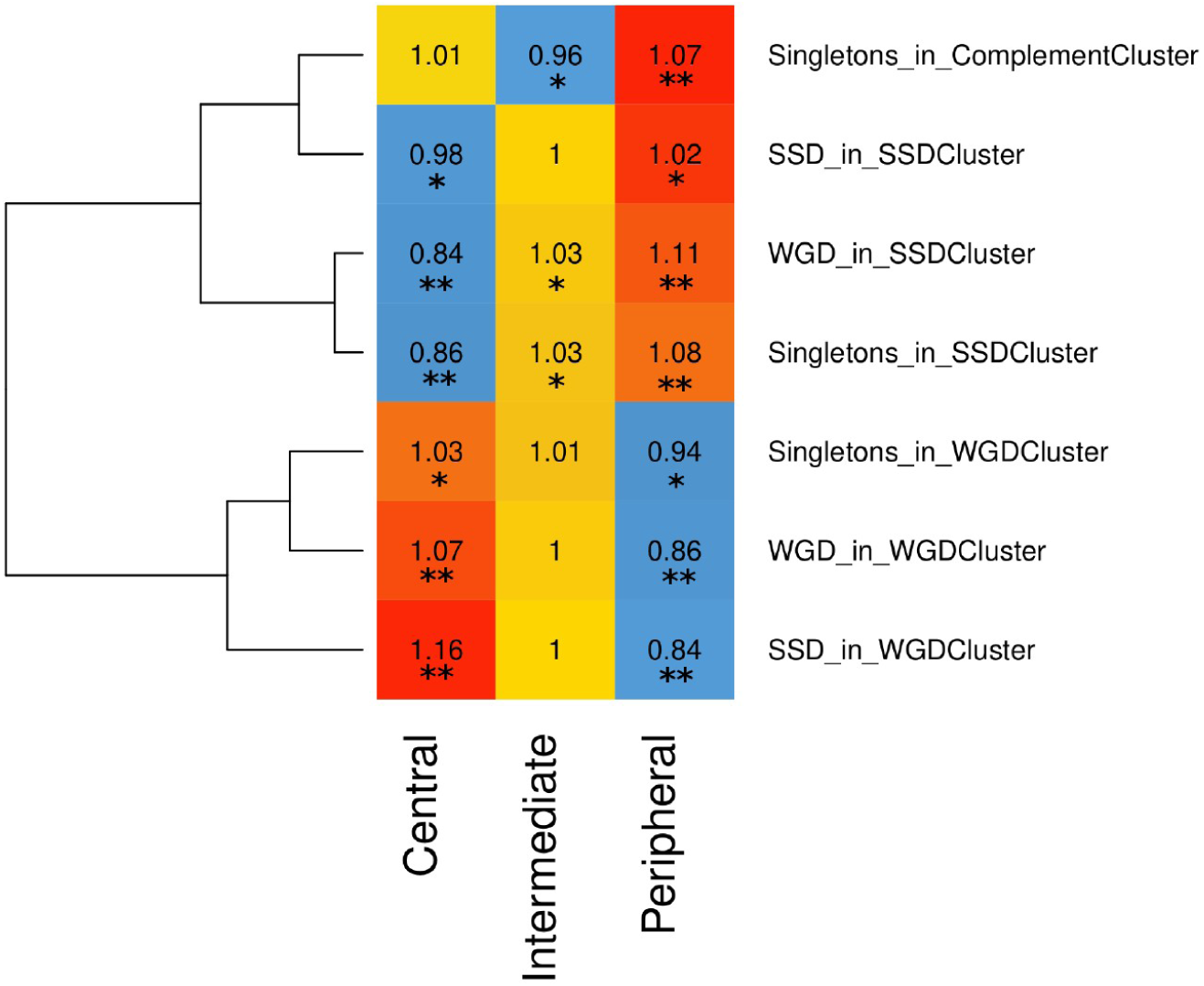
Relative enrichments of gene clusters in the three partitions of the yeast genome according to the polar distance from an assumed genome center (see text for details). Enrichments were calculated as observed over expected ratios of gene numbers, thus having a baseline value of 1. Significance assessed with permutation tests (**, p-value<=0.01; *, p-value <=0.05).

**Supplementary Figure 3.**
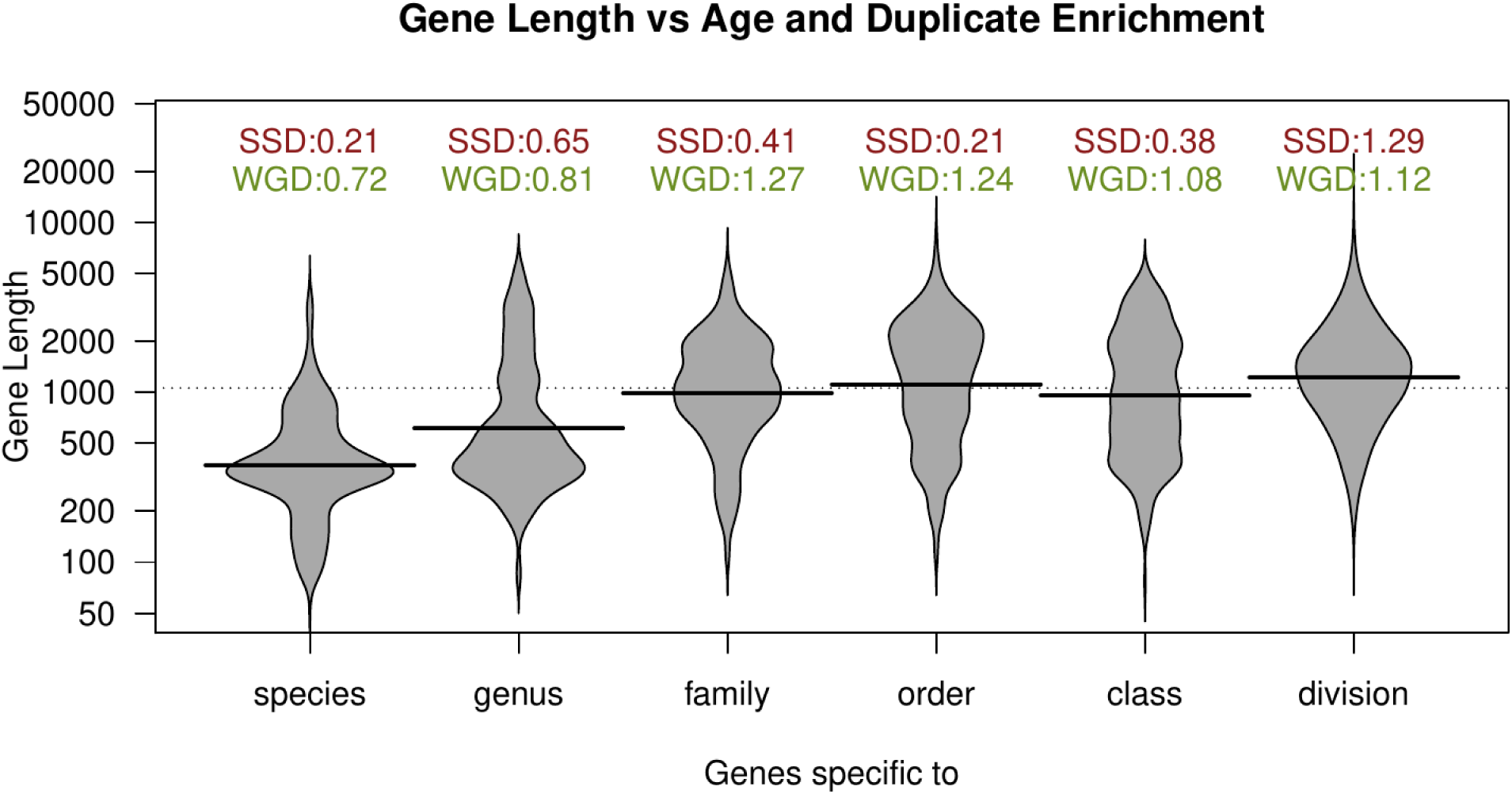
Gene length for sets of yeast genes that are specific to different age groups, accompanied by the relative enrichments of duplicate genes in each group (baseline enrichment is 1).

**Supplementary Figure 4.**
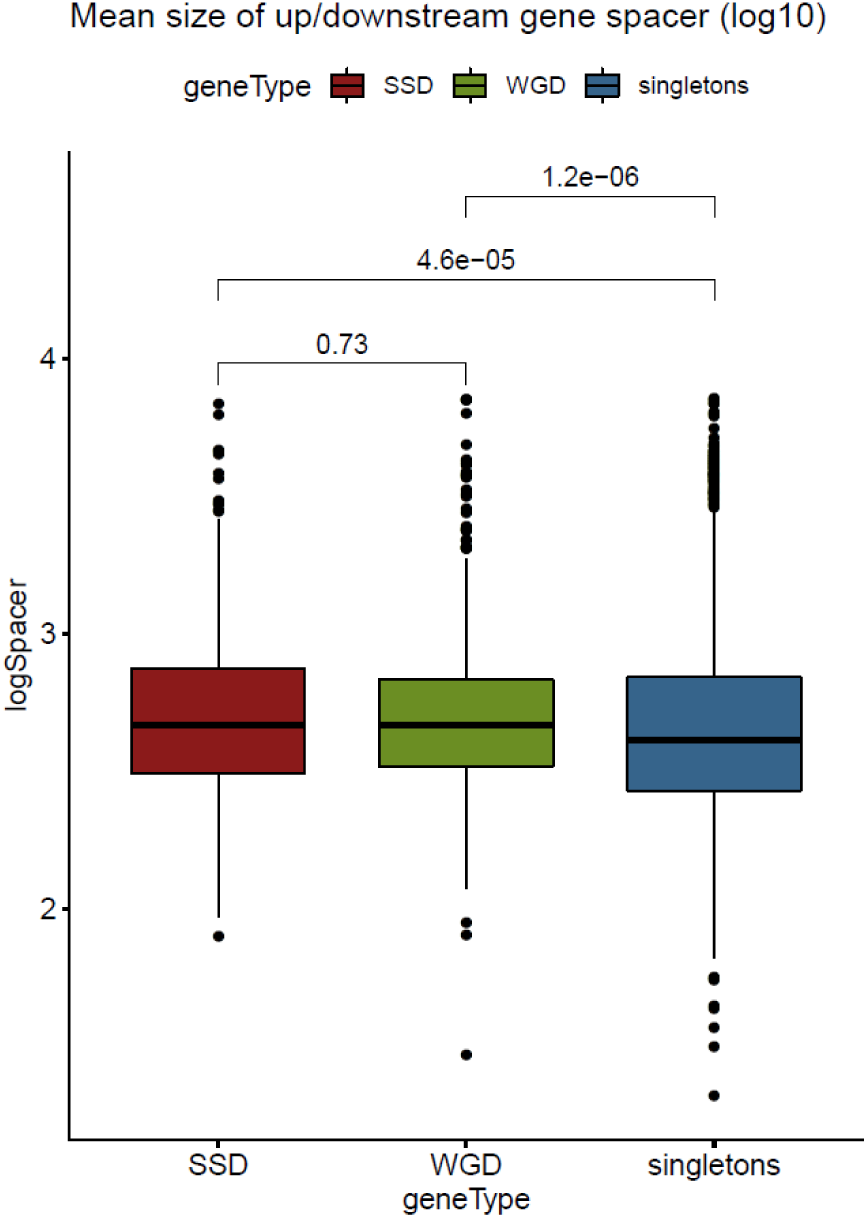
Distribution of the mean size of gene upstream and gene downstream distances of SSD, WGD and singleton genes form their flanking genes. Values over brackets denote p-values of a Mann Whitney test.

**Supplementary Figure 5.**
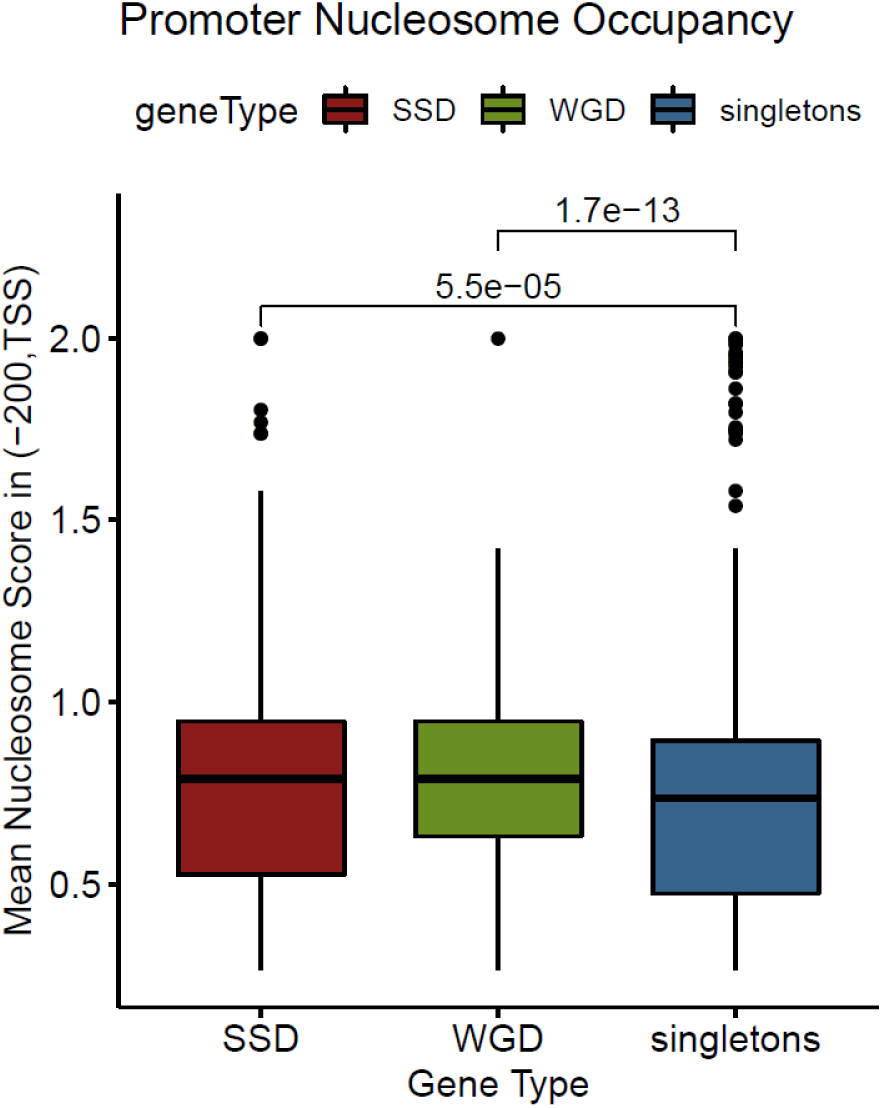
Distribution of mean nucleosome occupancy scores for a 200bp region immediately upstream of the TSS for SSD, WGD and 1000 random singleton genes. Values over brackets denote p-values of a Mann Whitney test.

**Supplementary Figure 6.**
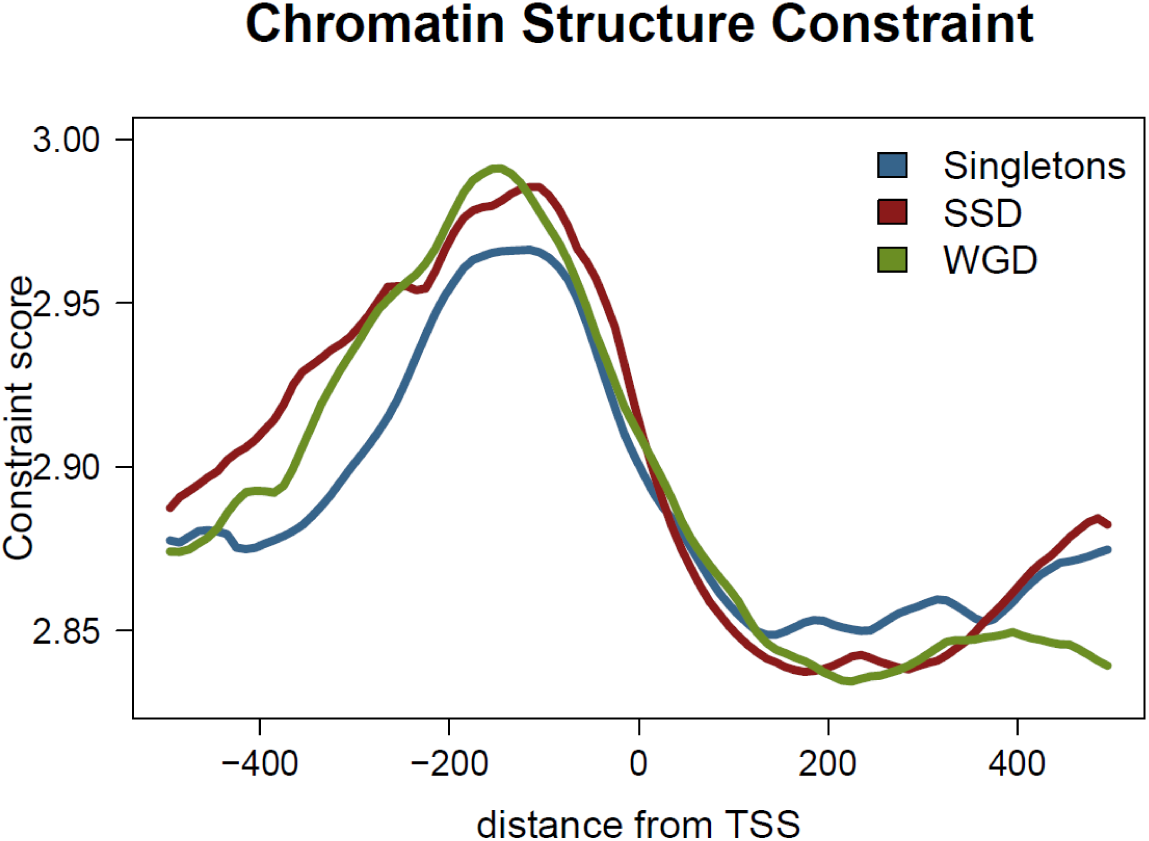
Chromatin structure constraint measured as mean SymCurv Robustness (Nikolaou et al. 2010) along a region spanning 500bps either side of TSS for singleton, SSD and WGD genes.

**Supplementary Figure 7.**
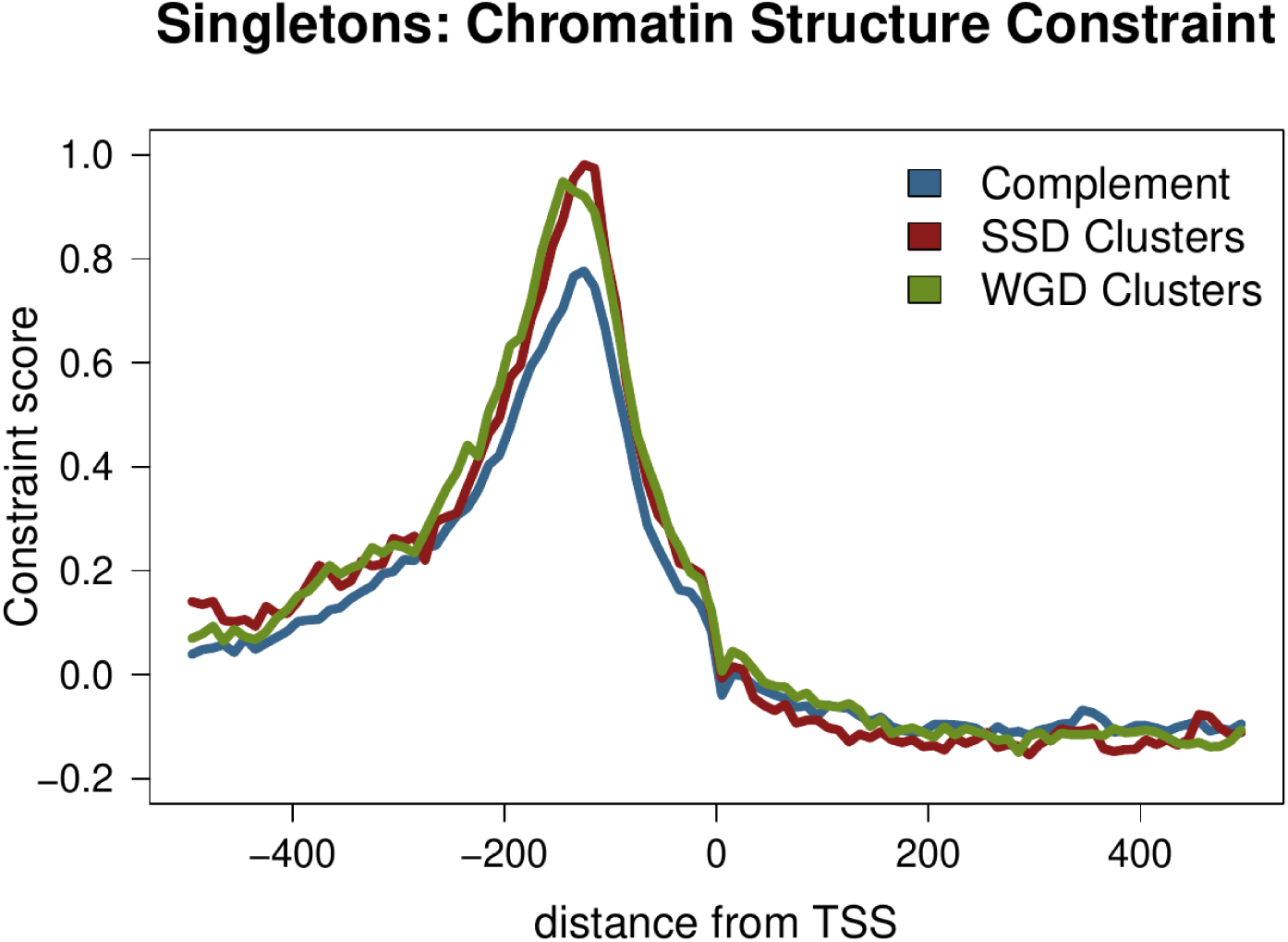
Chromatin structure constraint score as measured with the “mutation score” (Routhier et al, 2020) along a region spanning 500bps either side of TSS for singleton genes residing in the genome complement, SSD and WGD clusters.

**Supplementary Figure 8.**
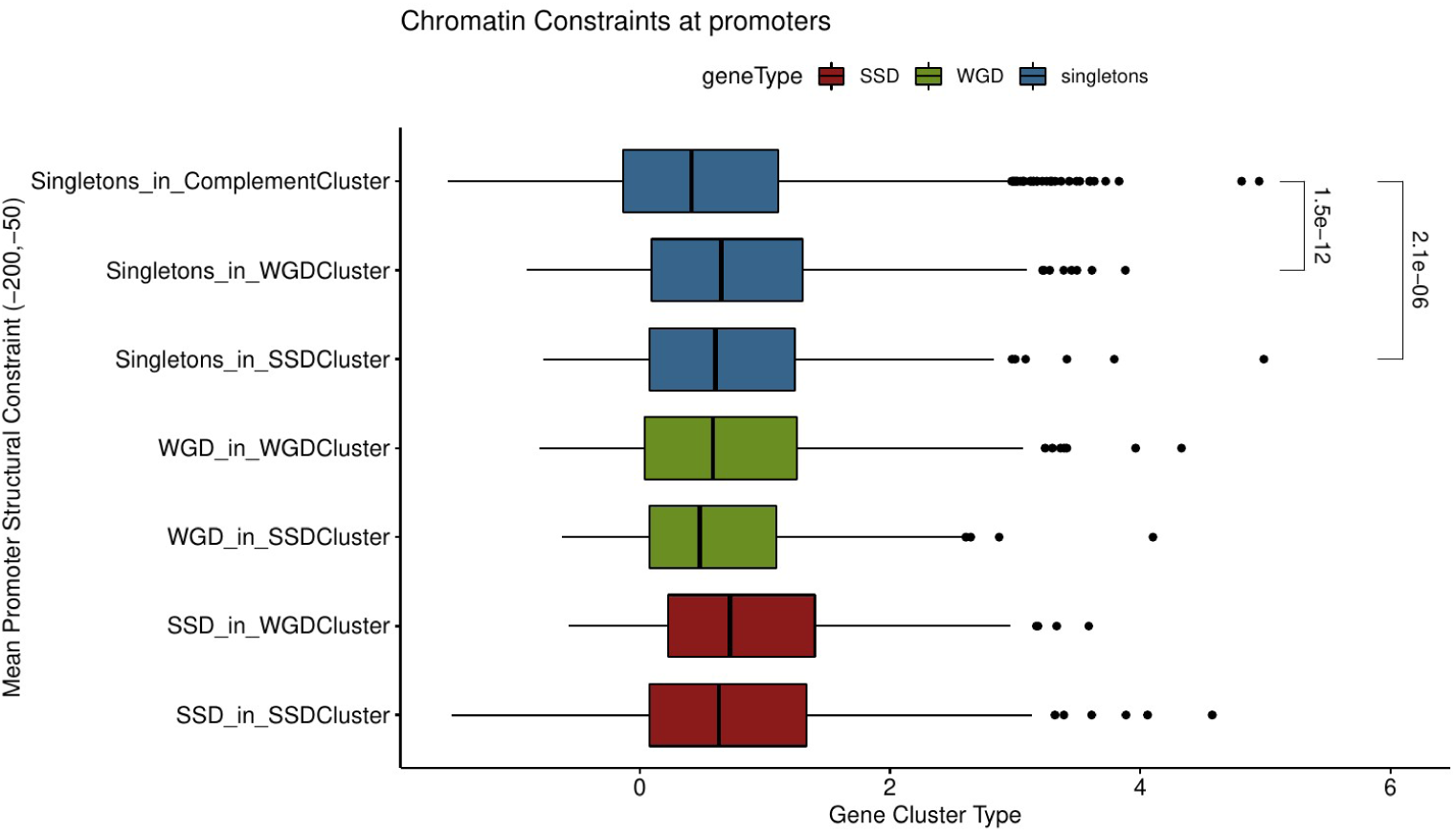
Distribution of mean promoter structural constraint measured as mutation score conservation in the proximal promoter region (200bp to 50bp upsteam to TSS) measured as mutation score (Routhier et al. 2020). Values next to brackets denote p-values of a Mann Whitney test.

**Supplementary Figure 9.**
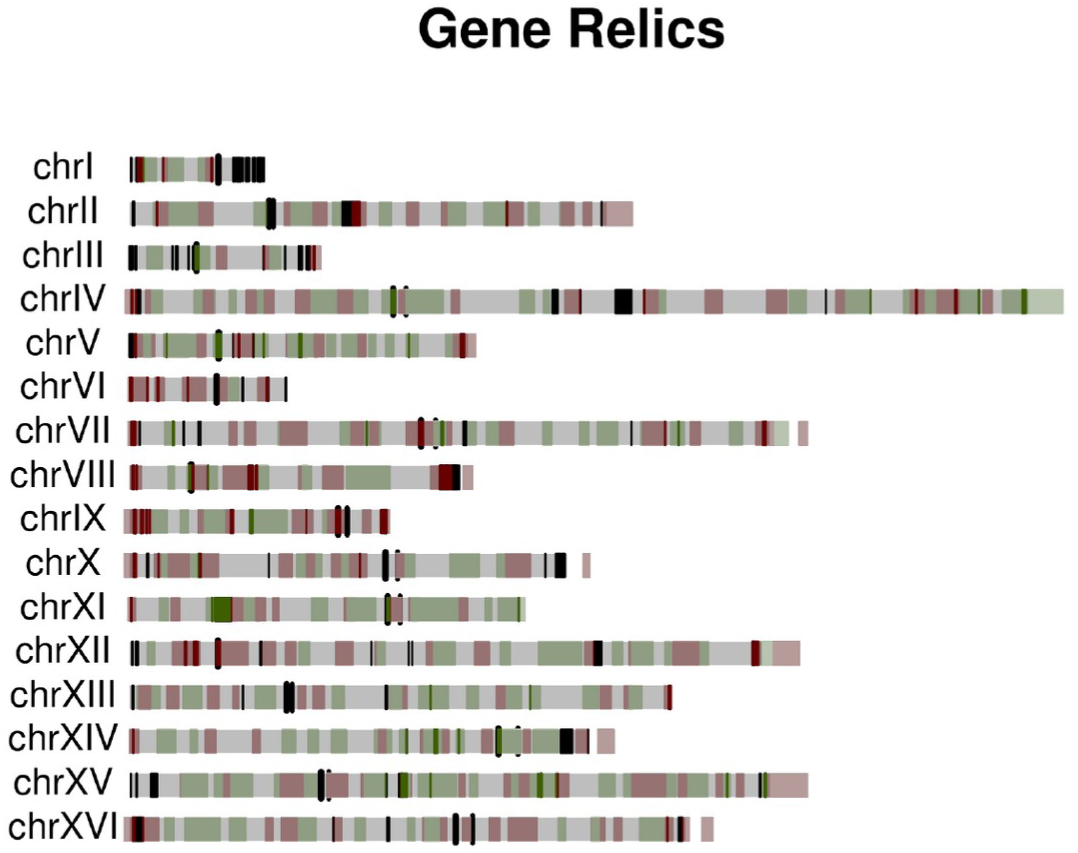
Gene relics (vertical bars) against the SSD (red) and WGD (green) gene clusters in the yeast genome.

**Supplementary Figure 10.**
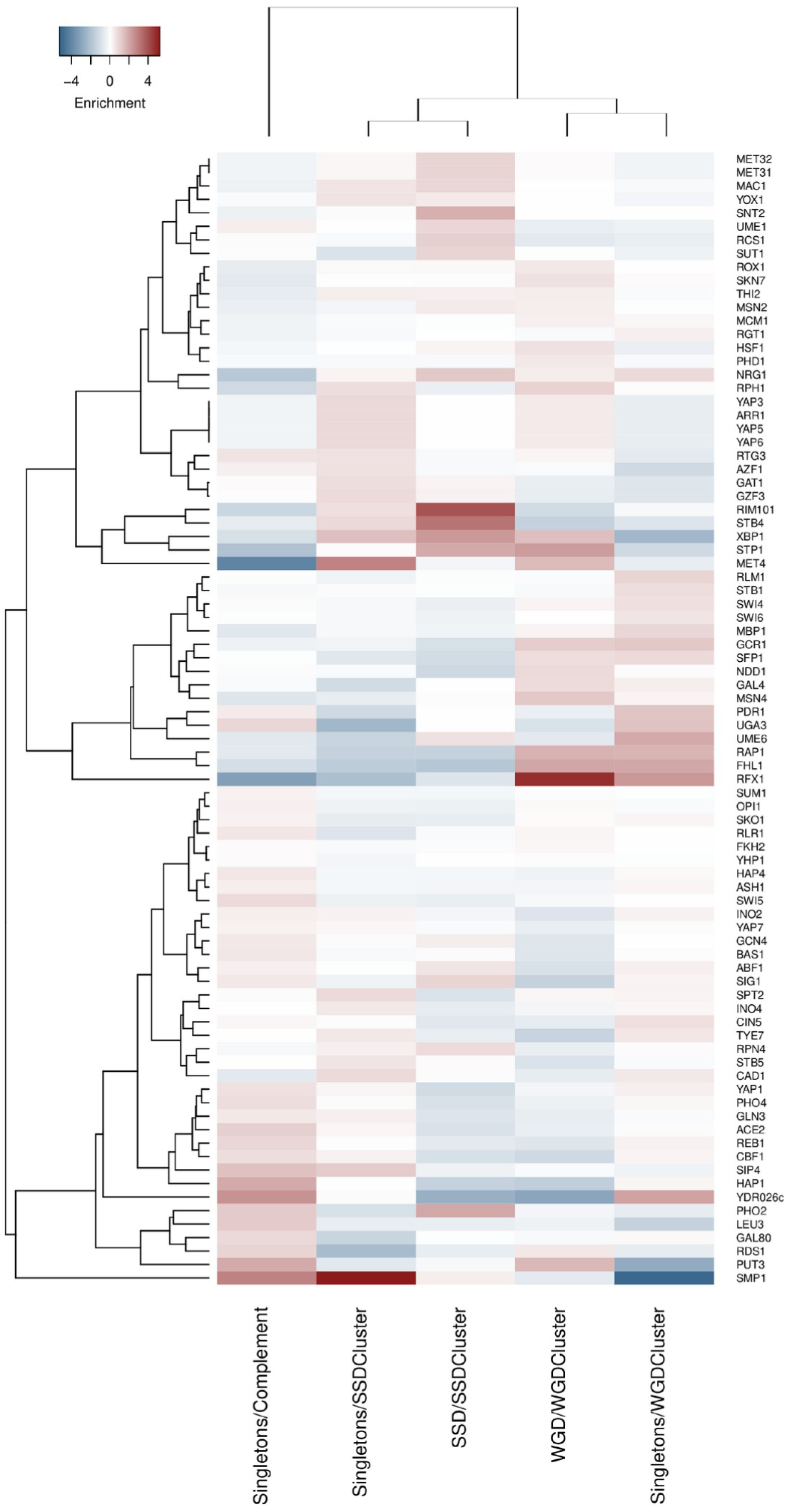
Enrichment heatmap of transcriptional regulator binding sites in different gene categories. Enrichments calculated as log2(observed/expected ratios).

**Supplementary Figure 11.**
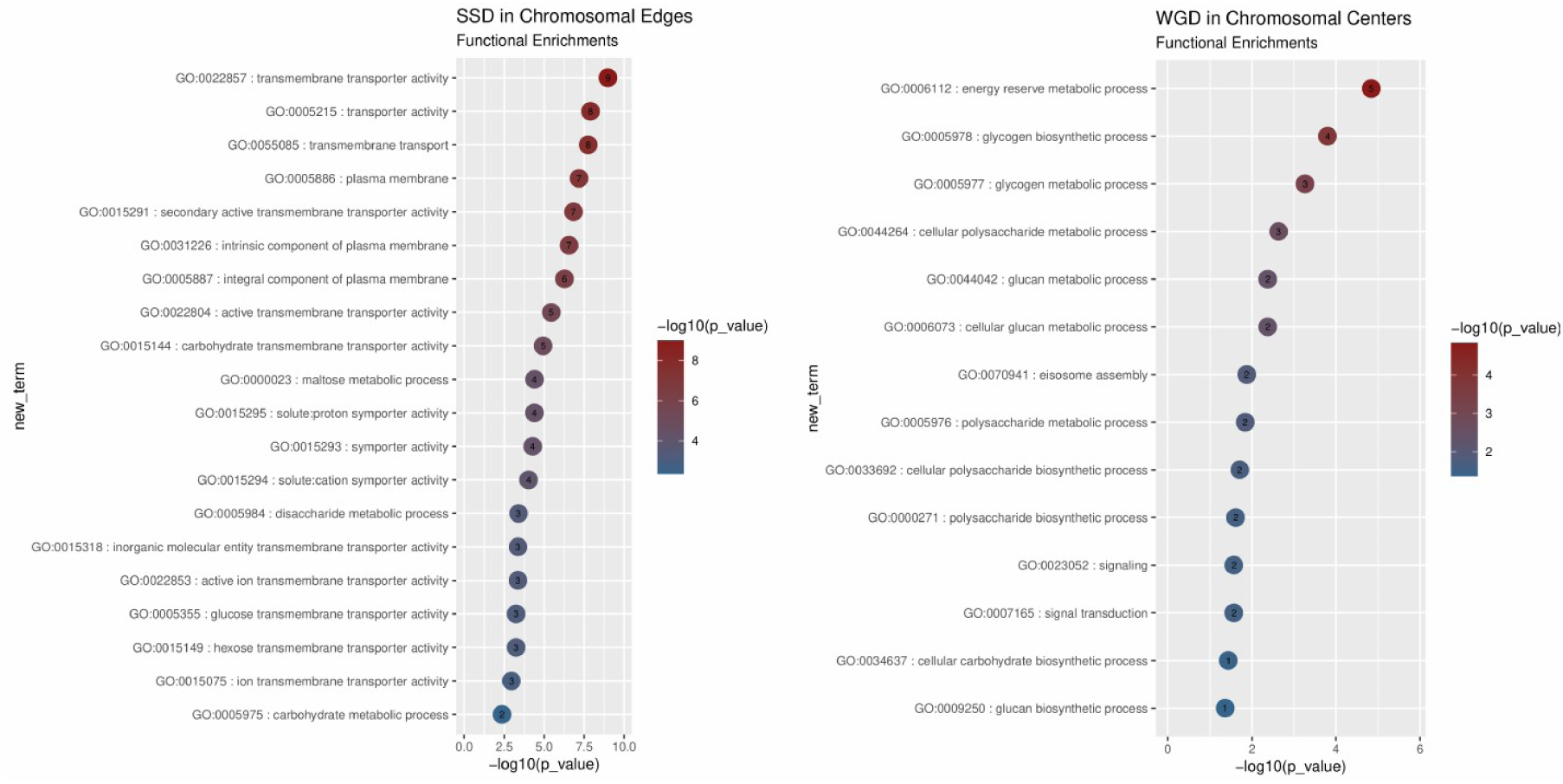
Functional enrichment plots representing the top most enriched GO terms for genes found in a) SSD genes in SSD clusters that lie close to the chromosomal edges (<10% of the chromosomal arm length) and b) WGD in WGD clusters found near the centromeres (>90% of the chromosomal arm length away from the edge).

